# Insights into the causes and consequences of DNA repeat expansions from 700,000 biobank participants

**DOI:** 10.1101/2024.11.25.625248

**Authors:** Margaux L.A. Hujoel, Robert E. Handsaker, Nolan Kamitaki, Ronen E. Mukamel, Simone Rubinacci, Pier F. Palamara, Steven A. McCarroll, Po-Ru Loh

## Abstract

Expansions and contractions of tandem DNA repeats are a source of genetic variation in human populations and in human tissues: some expanded repeats cause inherited disorders, and some are also somatically unstable. We analyzed DNA sequence data, derived from the blood cells of >700,000 participants in UK Biobank and the *All of Us* Research Program, and developed new computational approaches to recognize, measure and learn from DNA-repeat instability at 15 highly polymorphic CAG-repeat loci. We found that expansion and contraction rates varied widely across these 15 loci, even for alleles of the same length; repeats at different loci also exhibited widely variable relative propensities to mutate in the germline versus the blood. The high somatic instability of *TCF4* repeats enabled a genome-wide association analysis that identified seven loci at which inherited variants modulate *TCF4* repeat instability in blood cells. Three of the implicated loci contained genes (*MSH3*, *FAN1*, and *PMS2*) that also modulate Huntington’s disease age-at-onset as well as somatic instability of the *HTT* repeat in blood; however, the specific genetic variants and their effects (instability-increasing or-decreasing) appeared to be tissue-specific and repeat-specific, suggesting that somatic mutation in different tissues—or of different repeats in the same tissue—proceeds independently and under the control of substantially different genetic variation. Additional modifier loci included DNA damage response genes *ATAD5* and *GADD45A*. Analyzing DNA repeat expansions together with clinical data showed that inherited repeats in the 5’ UTR of the glutaminase (*GLS)* gene are associated with stage 5 chronic kidney disease (OR=14.0 [5.7–34.3]) and liver diseases (OR=3.0 [1.5–5.9]). These and other results point to the dynamics of DNA repeats in human populations and across the human lifespan.

## Introduction

Short tandem repeats (STRs) in which segments of 1–6 base pairs of DNA are repeated many times are mutable genomic elements with diverse influences on cellular and organismal phenotypes^1^. Common STR polymorphisms in human populations have been characterized using short-read^2,3^ and long-read DNA sequencing^4,5^ and shown to influence gene expression^6–8^ and complex traits^9,10^, while rare STR expansions are known to cause more than 60 genetic disorders, half of which were discovered in the past decade^11,12^. The allelic diversity that underlies these effects is generated by the high mutability of STRs: common STR alleles expand and contract in the germline orders of magnitude more frequently than single nucleotides mutate^13,14^, and rare, pathogenic STR expansions have long been observed to be unstable both intergenerationally and somatically^15^. Recently, genome-wide association studies (GWAS) have provided insights into the molecular mechanisms that modulate repeat instability^16^, both by directly ascertaining de novo STR mutations^17^ and by searching for genetic modifiers of the timing or progression of Huntington’s disease^18–23^, which are driven by somatic expansion of a CAG trinucleotide repeat in *HTT*^24^. These studies, so far of up to 16,399 persons with Huntington’s disease, have provided clues toward potential therapeutic targets for slowing or halting repeat expansion disorders.

Whole-genome sequencing (WGS) of biobank cohorts offers opportunities to identify new repeat expansion disorders and, in theory, to study germline and somatic instability of STRs in much larger sample sizes than previously possible. Here, we deeply analyzed repeat instability at 15 highly polymorphic CAG-repeat loci using available short-read WGS data from the blood-derived DNA of 490,416 participants in UK Biobank (UKB)^25,26^ and 245,388 participants in *All of Us* (AoU)^27^. To do so, we developed computational techniques for estimating the length and instability of DNA repeats from large numbers of short WGS reads^28^. These new methods enabled us to use data from very large numbers of research participants to learn about germline and somatic repeat instability, characterize allele-specific expansion and contraction rates of common repeat alleles, identify genetic influences on somatic repeat expansion, and discover pathogenic effects of expanded repeats.

## Results

### CAG trinucleotide repeat expansions in UK Biobank

We first sought to identify which CAG repeat loci in the human genome have expanded alleles present in the large UKB cohort (n=490,416). We identified UKB participants with long CAG repeat alleles (245 repeat units) by analyzing whole-genome sequencing data for the presence of 151bp sequencing reads comprised entirely or almost entirely of CAG repeat units (in-repeat reads (IRRs); Fig. 1A). Such reads were easily extractable from WGS read alignments previously generated using bwa^29^, as bwa aligned nearly all of these IRRs to the *TCF4* repeat sequence (which contains the longest CAG or CTG repeat tract in the GRCh38 human reference genome) even when they originated from loci other than *TCF4* (Supplementary Fig. 1). For each participant with one or more IRRs, we then determined the locus or loci from which the IRRs originated by ascertaining the mapping locations of the mate pairs of the IRRs, requiring mate sequences to map to loci that contain commonly-polymorphic CAG repeats^3^.

**Figure 1:**
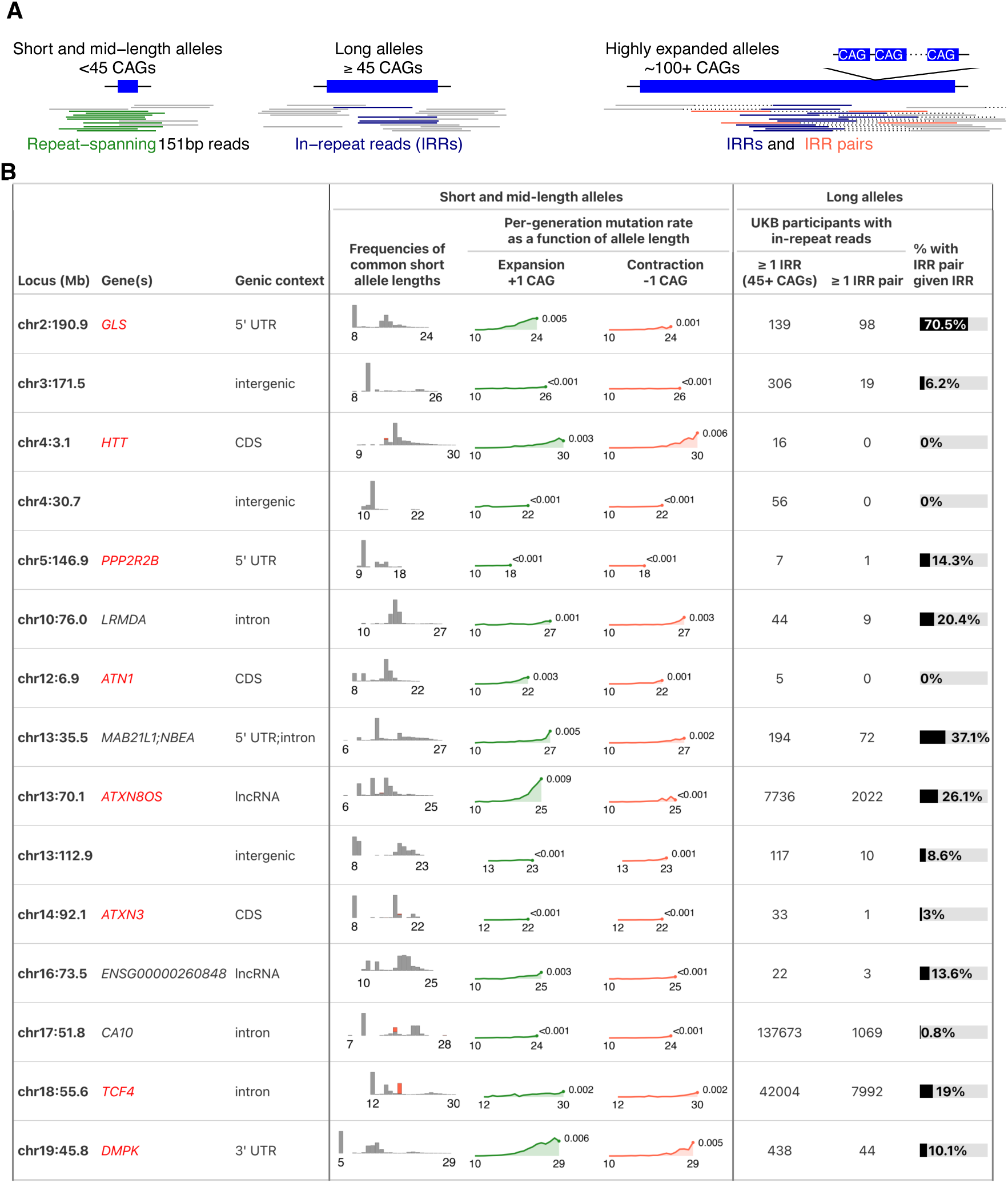
CAG trinucleotide repeat expansions in UK Biobank. **(A)** Types of short-read evidence of repeat alleles of different lengths. **(B)** Fifteen CAG repeat loci at which at least five UKB participants carried long alleles (≥45 repeat units). Repeat expansions in genes highlighted in red are known to be pathogenic. For each locus, the length distribution of common short alleles (≤30 repeat units) is shown; the length range is indicated below each histogram, and red bars denote interrupted repeat alleles. For each common allele between 10–30 repeat units, rates of intergenerational expansion and contraction (by ±1 unit) are plotted as a function of allele length; the mutation rate of the longest allele is indicated at the end of each curve. For long alleles, counts of UKB participants with at least one in-repeat read (IRR) or IRR pair are shown.

The vast majority of CAG repeat expansions in UKB occurred at only a few repeat loci: 18 autosomal CAG repeat sequences in the human genome were expanded to ≥45 repeat units in at least five UKB participants (Fig. 1B and Supplementary Table 1). Three repeat loci were expanded in thousands of UKB participants—*CA10* (137,673 participants), *TCF4* (42,004), and *ATXN8OS* (7,736)—together accounting for >97% of all observed expansions beyond 45 repeat units. Most of these repeats (15 of 18) were in transcribed regions of the genome, and exonic repeats appeared to be particularly enriched for long alleles (OR=3.19 [1.12–9.39], p=0.02; Supplementary Table 2), consistent with the idea that transcription contributes to repeat instability^15,30^. For nine of the 18 repeats, expanded alleles are known to be pathogenic^11^.

Given that ≥45-repeat-unit alleles of each of these repeats were observed in UKB, we reasoned that these repeat loci might comprise some of the most mutable CAG repeats in the human genome, such that analyzing all alleles of each repeat (including common, shorter alleles) in all UKB participants might provide insights into repeat instability. We therefore measured the lengths of short alleles of each repeat (≤30 repeat units) by analyzing sequencing reads that spanned the repeat, focusing on 15 of the 18 repeat loci that that passed further filters (Fig. 1 and Supplementary Table 1). These analyses recovered repeat length distributions consistent with previous analyses (Fig. 1B)^31,32^. In addition to allele length, we genotyped intra-repeat sequence variation (i.e., repeat “interruptions”), observing common interrupted alleles of four repeats (Fig. 1B).

### Germline instability of common CAG-repeat alleles

Short tandem repeats are a mutable class of genetic variation, generating ∼50–60 de novo repeat-length mutations per offspring^14,17,33^ among ∼1 million polymorphic STRs that exhibit a wide range of per-locus mutation rates^34,35^. Here, whole-genome sequencing of 490,416 UKB participants provided a unique opportunity to estimate allele-specific intergenerational expansion and contraction rates of each repeat locus. To do so, we analyzed length discordances among alleles belonging to genomic tracts inherited identical-by-descent (IBD) from shared ancestors, building upon IBD-based analyses of single-nucleotide mutations^36–38^ (Fig. 2A). Briefly, for each repeat, among individuals whose two alleles differed in length and could be confidently phased to SNP haplotypes, we identified each haplotype’s longest IBD partner (usually sharing >10 cM of IBD^39^). For each such IBD pair, we estimated the time to their most recent common ancestor (TMRCA), and we determined the allele carried by the common ancestor by examining alleles carried on “outgroup” haplotypes (Fig. 2A). Restricting to IBD pairs for which the ancestral allele was confidently determined, we obtained a data set containing hundreds of thousands of ancestral alleles together with the alleles transmitted to pairs of UKB participants (typically descended 10–30 generations; Supplementary Fig. 2A). The large number of allele transmissions represented in this data set—comprising millions of meioses—allowed us to precisely estimate allele-specific, expansion-or contraction-specific mutation rates.

**Figure 2:**
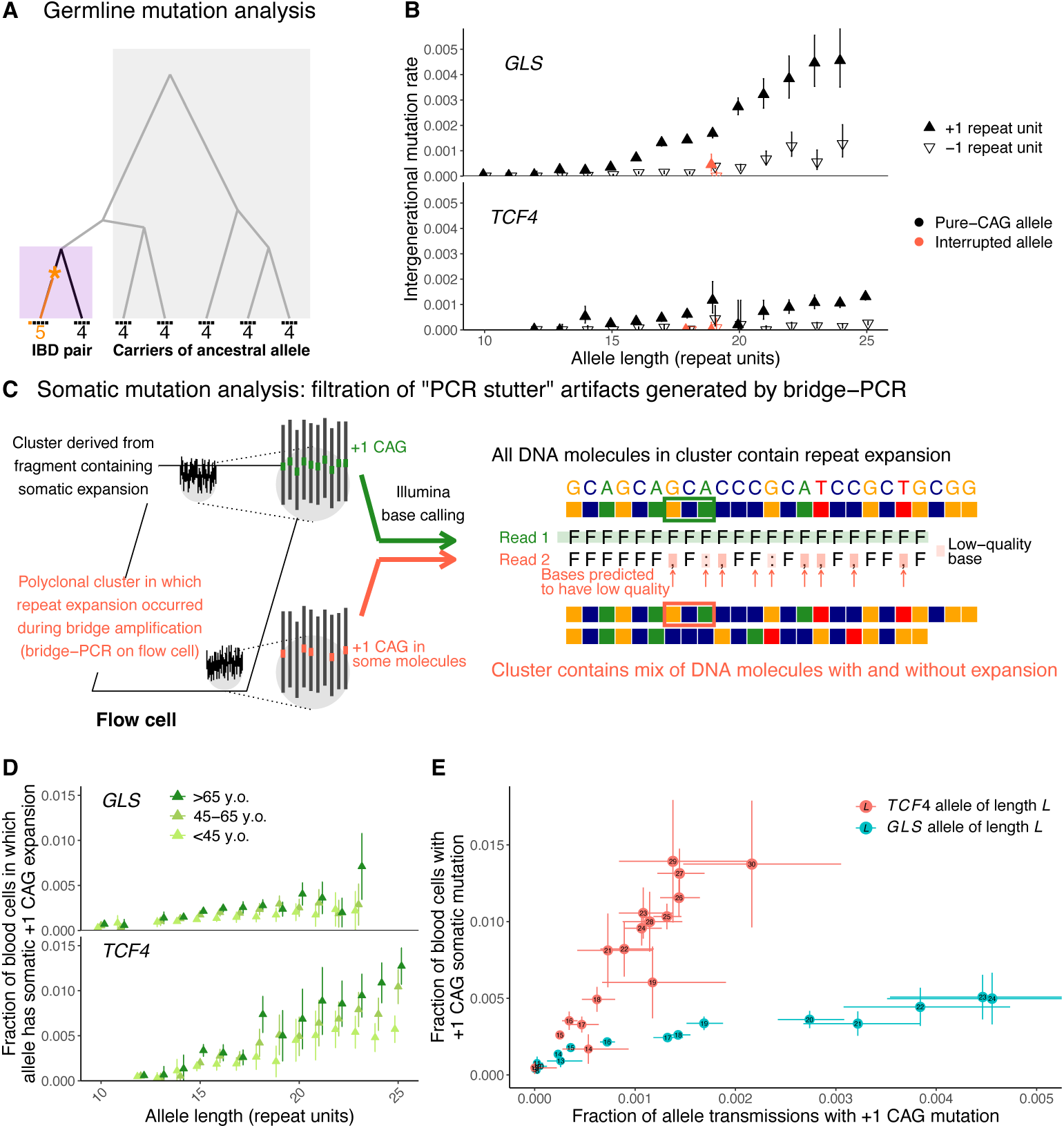
Germline and somatic instability of common CAG repeat alleles. **(A)** Germline mutation rates were estimated by analyzing discordance rates among alleles inherited within IBD tracts shared by pairs of UKB participants. Ancestral alleles were imputed from more-distantly shared haplotypes. **(B)** Per-generation rates of germline expansion (+1 repeat unit) and contraction (–1 repeat unit) of *GLS* and *TCF4* repeat alleles, estimated in UKB. **(C)** Analytical strategy for estimating somatic mutation rates by detecting and filtering out reads likely to reflect PCR artifacts introduced during sequencing. During (PCR-based) bridge amplification on a flow cell, a DNA fragment is clonally amplified into a cluster of colocalized DNA molecules. A PCR stutter error results in a polyclonal cluster containing a mixture of DNA molecules with and without the error. If the molecules containing the error constitute the majority of the cluster, the sequencing read generated from the cluster (reflecting the majority base at each position within the read) will contain the error, but the heterogeneity of the cluster will reduce base qualities at positions within the read that mismatch between molecules with and without the error. **(D)** Rates of somatic expansion of *GLS* and *TCF4* repeat alleles (i.e., fractions of blood cells in which an allele has expanded by +1 repeat unit), stratified by age in AoU. **(E)** Somatic mutation rates (in UKB) plotted against germline mutation rates for *GLS* and *TCF4* repeat alleles. Error bars, 95% CIs.

To validate this approach, we verified that estimated intergenerational mutation rates were consistent with rates of discordances between genotypes of sibling pairs (Supplementary Fig. 2B), and we also verified that probabilities of mutation between ancestral and transmitted alleles scaled linearly with estimated TMRCA (Supplementary Fig. 2A). We further verified that the distribution of mutational jump sizes obtained from this analysis was broadly consistent with distributions previously observed in analyses of de novo mutations^14,17^: across all 15 repeat loci, 61% of mutations modified CAG-repeat lengths by 1 repeat unit, with 44% being single-repeat-unit expansions and 17% being single-repeat-unit contractions. We focused further analyses on these most-common single-repeat-unit mutations.

Across all 15 repeat loci, intergenerational mutation rates increased with allele length, rising to 0.5–0.9% per generation for single-repeat-unit expansions of the longest common alleles of repeats in *GLS*, *DMPK*, and *ATXN8OS* (Fig. 1B and Supplementary Fig. 3). The mutation rates of these STRs far exceeded the genome-wide average, estimated to be ∼5×10^-5^ per haplotype per generation^14,17,33^. Repeat loci tended to either expand more often than contract (particularly so for *ATXN8OS* and *GLS*) or to have similar expansion and contraction rates (Fig. 1B and Supplementary Fig. 3). Single-nucleotide variants that interrupted repeat sequences greatly stabilized alleles: a common 18-repeat *TCF4* allele that contains an interruption in its ninth repeat unit exhibited a 135-fold [54–336] lower expansion rate compared to the uninterrupted 18-repeat allele, and an interruption in the second-to-last repeat unit of a 19-repeat *GLS* allele decreased intergenerational expansion rate 3.7-fold [1.9–7.2] (Fig. 2B). These results corroborate previous observations that repeat interruptions stabilize expansion of pathogenic alleles^12,40–43^ and quantify the strength of such effects in the germline.

### Somatic expansion of common CAG-repeat alleles in blood

These high rates of germline instability led us to wonder whether common alleles of some repeats might be sufficiently unstable in blood cells for somatic length-change mutations to be observable in short-read WGS data. Somatic expansions and contractions of STRs are challenging to identify from WGS data because polymerase slippage during PCR amplification can spuriously alter repeat lengths^44–46^. Such “PCR stutter” errors arise even when sequencing libraries are prepared using a PCR-free protocol because PCR still occurs during the bridge amplification step of Illumina sequencing by synthesis^47^. However, we realized that this PCR error mode tends to produce predictable “barcode” patterns of reduced base quality scores within sequencing reads, thereby allowing most artefactual reads to be detected and excluded from analysis (Fig. 2C and Supplementary Fig. 4A) and enabling detection of mutations present in smaller proportions of cells than could previously be recognized.

We applied this filtering strategy to estimate repeat-specific, allele-specific somatic expansion rates in UKB, which we quantified as the average fraction of blood cells in which a given repeat allele has expanded by 1 repeat unit. For each common repeat allele, we estimated this quantity by computing the average number of reads containing a single-repeat-unit expansion of the allele (in individuals heterozygous for the allele) and normalizing by the average number of reads observed in individuals heterozygous for the one-repeat-longer allele (after imposing the same filters on base qualities). The age range of the UKB cohort (40–70 years of age at DNA acquisition) enabled us to assess the efficacy of the filtering strategy: if filtering worked perfectly, the fraction of blood cells harboring a somatic mutation should increase approximately linearly with age, whereas if filtering worked poorly (such that most putatively somatic mutations were actually technical artifacts), the fraction of blood cells estimated to harbor a somatic mutation should have no relationship with age.

For four of the 15 repeats (in *TCF4*, *GLS*, *DMPK*, and *ATN1*), we detected significant increases in somatic single-repeat-unit expansion rates with age, suggesting that common alleles of these repeats expand with sufficient frequency in blood cells for mutations to be detectable above residual PCR error (Supplementary Fig. 5). These findings replicated in the AoU data set, in which the wider age range of participants (18 to ≥90 years of age at blood draw) produced clear increases in fractions of blood cells harboring somatic single-repeat-unit expansions in older versus younger individuals (Fig. 2D and Supplementary Fig. 6). Somatic instability in blood increased with allele length for all four repeats (Fig. 2D and Supplementary Fig. 6), such that for longer common alleles, the large majority of putatively somatic single-repeat-unit expansions appeared to reflect real somatic mutations rather than technical artifacts (Supplementary Fig. 4B,C). Repeat alleles in *TCF4* were the most somatically unstable: in carriers of alleles containing 25 or more repeat units, we estimated that at least 1% of blood cells contained a somatic expansion by the time an individual reached age 55 (Fig. 2D), such that detectable mosaicism at *TCF4* is common by middle age. In contrast, we did not detect evidence of age-associated contraction of any of the 15 repeat loci, probably owing to a combination of lower somatic contraction rates and higher rates of residual PCR stutter errors, which tend to be contraction-biased^44^.

Comparing these estimates of somatic one-repeat-unit expansion rates with our estimates of intergenerational mutation rates showed that the relative (blood/germline) rates of CAG-repeat expansion varied several-fold across repeat loci (Fig. 2E). While the *TCF4* repeat exhibited the greatest somatic instability in blood, it was relatively stable in the germline, whereas the *GLS* repeat displayed the opposite behavior (Fig. 2E), as did the *DMPK* repeat (Fig. 1B and Supplementary Figures 3 and 6). These results align with observations that somatic instability of pathogenic repeat expansions is highly tissue-specific, perhaps because of differences in transcription or trans-acting factors^30,48–52^. Consistent with the former hypothesis, the four repeats for which we detected instability in blood are in genes with significantly higher expression in blood (Wilcoxon rank-sum p=0.034; Supplementary Fig. 7).

### Somatic expansion of long *TCF4* alleles in blood

The high somatic expansion rates of *TCF4* repeat alleles—even those of shorter lengths— suggested the possibility that long *TCF4* alleles (245 repeat units) might be sufficiently unstable in blood to allow individual-level phenotyping of somatic expansion using short-read WGS data. If so, this would provide a unique opportunity to learn about instability of long repeats from somatic expansions in very many people—potentially enabling discovery of genetic modifiers of repeat instability^16,18–23,53^—as long *TCF4* alleles are unusually common (EUR AF=4%; 41,580 carriers in UKB; Fig. 1B).

A barrier to analyzing repeat expansions from widely available short-read WGS data is that ≥45-repeat-unit alleles are almost always too long to be spanned by short sequencing reads. It is not possible to directly measure lengths of alleles that exceed the length of a sequencing read (151bp), nor is it possible to directly observe mosaicism of alleles that have undergone varying amounts of somatic expansion. However, short-read WGS data does permit rough estimation of the length of a long allele by counting reads originating from the repeat: the longer an allele, the more in-repeat reads (IRRs) it should generate^54^. In an individual who is mosaic for somatic expansions of varying extents, this approach estimates the average length of expanded alleles across cells.

Applying this approach to UKB revealed a population-level pattern of somatic expansion of *TCF4* alleles with age: on average, lengths of ≥45-repeat-unit alleles increased with age of UKB participants (p=1.5×10^-116^), with mean allele lengths increasing from 81.9 (s.e.=0.3) repeat units in 40–44 year-olds to 92.6 (0.4) repeat units in 65–69 year-olds. However, these estimates were only weakly informative of individual-level somatic expansion, as they also reflected (i) variability in the lengths of long alleles inherited by different individuals; and (ii) considerable stochasticity in counts of IRRs (Supplementary Fig. 8A).

To facilitate further, better-powered analyses of somatic expansion, we devised two strategies to address these challenges. First, to control for variation in lengths of inherited *TCF4* alleles, we imputed the expected length of each individual’s long allele(s) from other UKB participants sharing a recent common ancestor. This allowed us to calibrate each individual’s measured allele length against measurements from other individuals sharing the same inherited allele (in lieu of another measurement in the same individual at a different time point). Suggesting that these imputation-based estimates were accurate, we found that controlling for imputed allele length strengthened the association of estimated allele length with age (p=6.5×10^-138^). Stratifying individuals by imputed allele length showed that somatic expansion accelerates rapidly with allele size, reaching ∼1 repeat unit per year for 100-repeat *TCF4* alleles (Fig. 3A).

**Figure 3:**
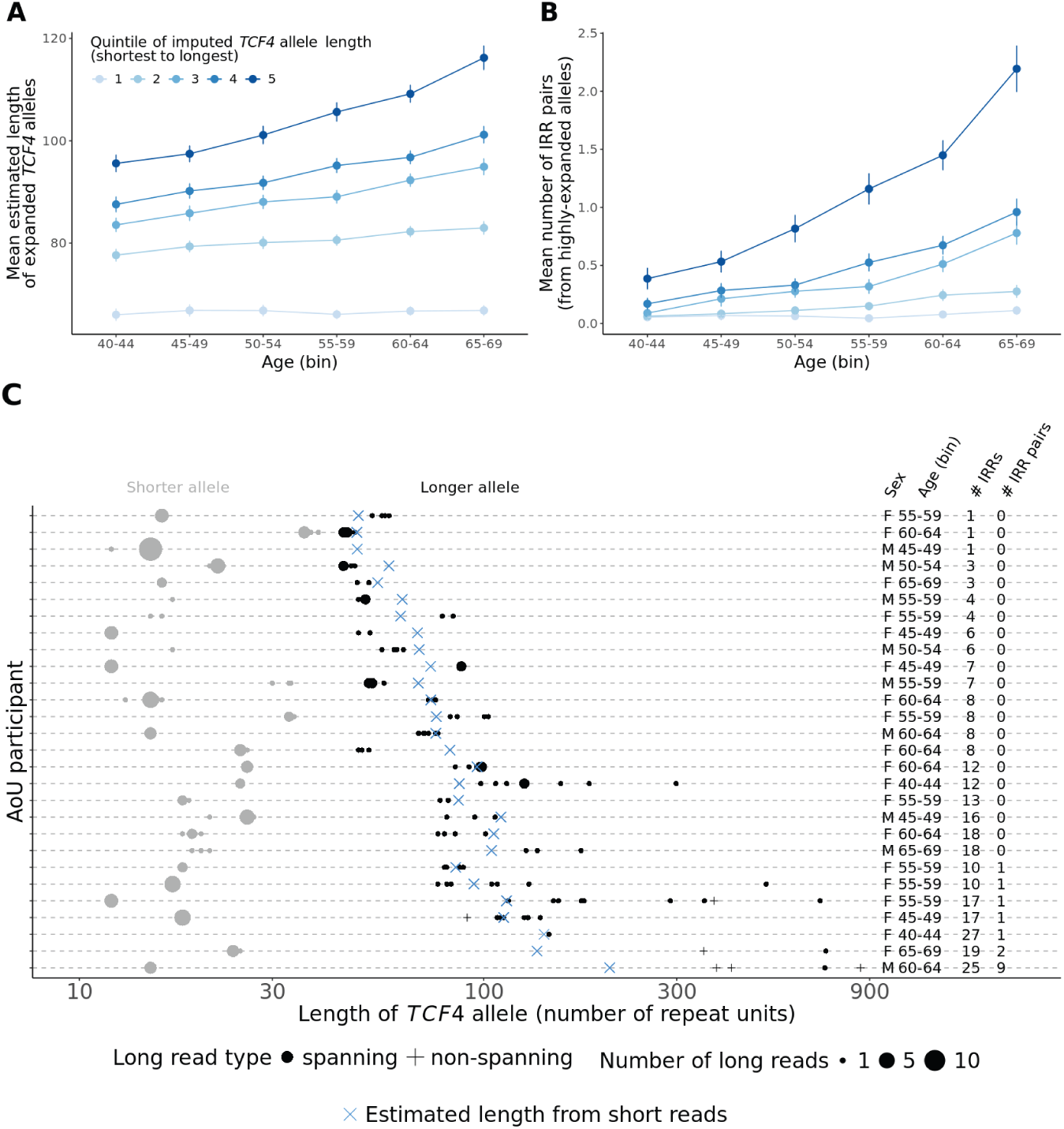
Somatic instability of long *TCF4* repeat alleles. **(A)** Mean estimated length (in repeat units) of long *TCF4* alleles (≥45 repeat units) in UKB participants of different ages. Heterozygous carriers of long *TCF4* alleles were first stratified into quintiles of imputed *TCF4* allele length, a proxy for inherited allele length. **(B)** Mean number of IRR pairs observed per UKB participant heterozygous for a long *TCF4* allele, again stratified by imputed *TCF4* allele length and by age. Analyses were restricted to individuals carrying no other long CAG repeat except possibly in *CA10* (such that IRR pairs could be assumed to have originated from *TCF4*). Error bars, 95% CIs. **(C)** *TCF4* allele lengths directly measured from long-read sequencing of carriers of long alleles in AoU. Each horizontal line corresponds to a single AoU participant; black markers indicate repeat lengths observed in long reads that span the *TCF4* repeat (dots) or partially overlap the repeat (pluses, which lower-bound allele lengths), while blue crosses indicate allele lengths estimated from short-read WGS. Long *TCF4* alleles exhibit somatic mosaicism, with alleles sometimes varying in length by hundreds of repeat units within blood cells from the same individual, indicating high somatic instability. We have received an exception from the *All of Us* Resource Access Board to disseminate participants counts less than 20.

Second, to reduce noise in estimates of long *TCF4* allele lengths, we computed an alternative metric based on the number of sequenced DNA fragments derived from a highly expanded repeat. Such fragments, which manifest in WGS data as in-repeat read pairs (IRR pairs; Fig. 1A), begin to be observed in 151bp paired-end sequencing data when alleles expand to ∼100 repeat units. The specificity of such read pairs to highly expanded alleles thus reduces the effect of sampling noise on estimates of allele lengths in this most-unstable length range (Supplementary Fig. 8A,B). The IRR pair metric associated even more strongly with age (p=1.3×10^-235^, controlling for imputed allele length and capping the count at 5), and among individuals in the top two quintiles of imputed allele length, mean counts of IRR pairs increased several-fold across the age range of UKB participants (Fig. 3B).

Long-read sequencing of blood-derived DNA from AoU participants (available for n=1,027 participants, of whom 28 had long *TCF4* alleles) corroborated *TCF4* allele length estimates from short-read WGS and demonstrated extensive mosaicism of somatically expanded alleles within individuals (Fig. 3C). Among individuals with highly expanded alleles (based on observing at least one IRR pair), every observed long read indicated a different allele length, and the range of observed allele lengths typically spanned hundreds of repeat units (Fig. 3C), consistent with recent observations^55,56^.

### Inherited genetic modifiers of somatic *TCF4* DNA-repeat expansion in blood

We used this common somatic-repeat-expansion phenotype to map common genetic influences on somatic repeat expansion. Studies of Huntington’s disease (HD) age-at-onset have identified several loci at which common genetic variants influence the age at which HD motor symptoms commence; many of these loci contain DNA-repair genes (such as *FAN1* and *MSH3*) that also affect the stability of DNA repeats^18–22^. The DNA repeat that causes HD appears to become toxic to striatal neurons only after considerable somatic expansion (to >150 CAG repeat units, generally from an inherited length <60 units), suggesting a likely reason that genetic modifiers tend to be in or near DNA-repair genes^24^.

To explore whether similar or different genetic effects might modulate repeat expansion in blood, we first examined the lead HD-associated SNP at *FAN1* (rs35811129) for association with *TCF4* expansion. This SNP indeed associated with estimated length of long *TCF4* alleles (p=1.3×10^-5^, adjusting for age, sex, and 20 PCs), and the association strengthened upon conditioning on imputed allele length (p=4.5×10^-6^) and using the alternative IRR pair-based length metric (p=1.1×10^-10^), mirroring the behavior of associations of these metrics with age. This suggested the possibility of further increasing GWAS power by constructing a *TCF4* somatic-expansion phenotype that more-optimally consolidates information from (i) IRR pairs and (ii) imputed allele lengths by using these two quantities to predict an individual’s age (the intuition being that repeat-expansion-predicted age might approximately capture an individual’s expected genetic liability for repeat expansion). Constructing a somatic-expansion phenotype in this way (Supplementary Fig. 8C) and incorporating IRR pairs observed in exome-sequencing data further strengthened association with rs35811129 (p=9.1×10^-13^).

Performing a GWAS on this *TCF4* somatic-expansion phenotype in UKB (n=40,231 participants satisfying sample inclusion criteria) identified four loci at genome-wide significance (p<5×10^-8^) (excluding *TCF4* itself, at which associations appeared to reflect imperfect control for inherited allele lengths; Supplementary Fig. 9) and five additional loci reaching suggestive significance (p<1×10^-6^; Supplementary Table 3). Three of the five suggestive signals replicated in an analogous analysis of AoU (n=8,217), indicating robustness of the association approach (Supplementary Table 3).

We therefore meta-analyzed GWAS results across UKB and AoU, identifying seven loci at which common inherited variants modulate *TCF4* repeat expansion (at p<5×10^-8^; Fig. 4A, Table 1, and Supplementary Fig. 10). Three loci—*MSH3* (p=2.0×10^-52^), *FAN1* (p=8.5×10^-29^), and *PMS2* (p=3.0×10^-8^)—overlapped mismatch repair-related genes known to modulate HD age-at-onset, and relative association strengths at these loci closely matched associations with the age-at-SDMT30 (symbol digit modalities test score 30) cognitive landmark in HD^21^ (Fig. 4A). The four other modifier loci included two DNA damage response genes: *ATAD5* (p=4.9×10^-12^), which was recently implicated in somatic expansion of *HTT* in blood^22^, and *GADD45A* (p=2.9×10^-8^), which encodes a growth arrest and DNA damage protein that binds R-loops^57^. Additionally, at 14q13.3 (near *SFTA3* and *NKX2-1*), a variant that associated with increased *TCF4* expansion appeared to also associate with delaying of the SDMT30 cognitive phenotype (p=0.013; Table 1).

**Figure 4:**
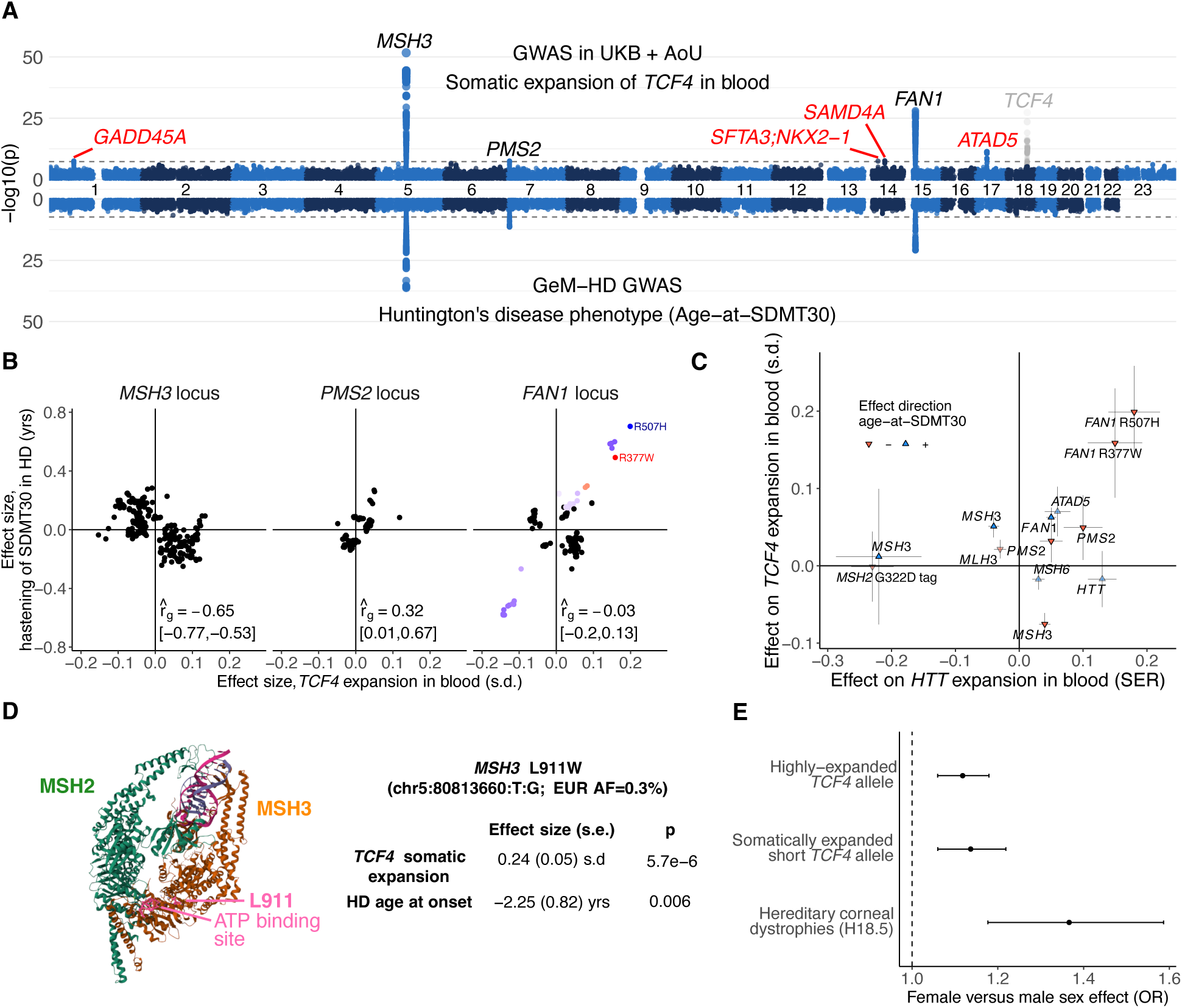
Genetic influences on somatic expansion of *TCF4* repeat alleles in blood. **(A)** Genome-wide associations with somatic instability of long *TCF4* repeat alleles in blood (top, meta-analyzed across UKB and AoU) compared to genetic associations with age-at-SDMT30, a Huntington’s disease clinical landmark of cognitive decline (bottom, from ref. ^21^). **(B)** Comparison of effect sizes of variants at *MSH3*, *PMS2*, and *FAN1* for hastening of SDMT30 in HD versus somatic expansion of *TCF4* repeats in blood. Local genetic correlation estimates at each locus are indicated. **(C)** Comparison of effect sizes for *TCF4* expansion in blood versus *HTT* expansion in blood (from ref. ^22^). **(D)** Effect sizes of a rare missense variant in *MSH3* (rs41545019, L911W) on *TCF4* expansion and HD age-at-onset (from ref. ^21^). The affected amino acid is indicated on a crystal structure of MutSβ complexed with a DNA insertion-deletion loop^69^ (PDB structure 3THZ). **(E)** Increased odds in female versus male UKB participants for having a highly-expanded *TCF4* allele (i.e., ≥1 IRR pair), a somatic expansion of a short *TCF4* allele (observed in a spanning short read), or a hereditary corneal dystrophy (ICD-10 code H18.5, which includes FECD). Odds ratios are from logistic regression adjusting for age, age squared and 20 PCs; for the association with presence of a highly-expanded allele, we also adjusted for the additional covariates included in our GWAS for somatic instability of long *TCF4* alleles. Error bars, 95% CIs.

**Table 1:**
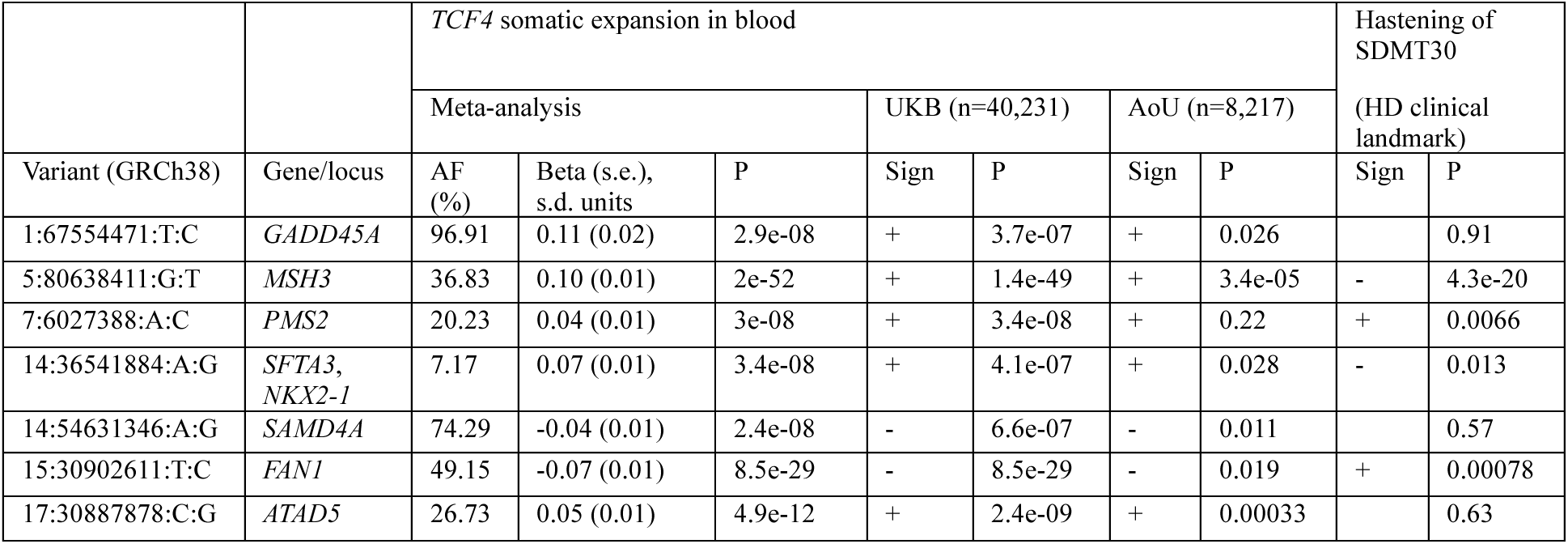
Genetic associations with somatic expansion of *TCF4* repeat alleles in blood. Allele frequencies, effect sizes (beta), and p-values are reported for lead variants from loci that reached genome-wide significance (p<5×10^-8^) in GWAS meta-analysis of somatic instability of long *TCF4* alleles in UKB and AoU (restricting to long-allele carriers with age at least 40). Effect directions and p-values within each cohort (UKB and AoU) are provided, and for comparison, p-values for hastening of SDMT30 (a clinical landmark of cognitive decline in Huntington’s disease^21^) are also provided. Effect sizes for hastening of SDMT30 are provided for associations with p<0.05.

Comparing these genetic modifiers of *TCF4* repeat expansion in blood to genetic modifiers of HD age-at-onset (which are likely regulating *HTT* repeat expansion in neurons^24^) revealed surprising and important differences. Counterintuitively, at both *MSH3* and *FAN1*, common haplotypes that decreased expansion of the *TCF4* repeat in blood appeared to increase expansion of the *HTT* repeat in the brain, based on associations with earlier onset of HD clinical symptoms^21^ (Fig. 4B, Supplementary Fig. 10A, and Supplementary Table 4). In contrast, two missense variants that reduce FAN1 activity^58^ appeared to increase expansion of both the *TCF4* repeat (in blood) and *HTT* repeat (in brain), as did common variants at *PMS2* (Fig. 4B). At each locus, the causal alleles (or the relative effect sizes of such alleles) appeared to be at least somewhat distinct across the two somatic-expansion settings (based on local genetic correlations 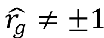; Fig. 4B); however, larger association studies will be needed to fine-map these genetic associations to causal variants.

Genetic modifiers also appeared to have different effects on expansion at different genomic loci in the same tissue. Among 13 variants recently observed to associate with *HTT* repeat expansion in blood^22^, 10 of the 13 also appeared to influence *TCF4* repeat expansion (p<0.05); however, only 6 of 10 had the same effect direction, with opposing effects observed at *MSH3*, *MLH3*, and *MSH6* (Fig. 4C and Supplementary Table 5). Moreover, the strongest modifier of *HTT* expansion in blood—a haplotype containing a missense variant in *MSH2* also implicated in germline STR mutation^17^—appeared not to affect *TCF4* expansion (p=0.54; Fig. 4C and Supplementary Table 5). These results suggest that the tissue-specific instability of many trinucleotide repeats^48–52^ may arise from complex regulation of mismatch repair processes that differs across cell types^22^ and even across repeat loci, perhaps interacting with locus-specific differences in chromatin structure or other epigenomic properties.

Follow-up analysis of protein-coding variation in loci influencing *TCF4* repeat expansion identified a secondary association of a low-frequency missense variant in *MSH3* (rs41545019; AF=0.3%, CADD score 28.1) with a large increase in *TCF4* instability (0.24 [0.05] s.d.; p=5.7×10^-6^), similar in magnitude to effects of the two missense variants in *FAN1* (Fig. 4D). This variant appeared to have a previously-undiscovered onset-hastening effect on HD (–2.25 (0.82) years; p=0.006 in ref.^21^). We speculate that the leucine-to-tryptophan substitution encoded by rs41545019 (L911W, which is near a conserved ATP binding site; Fig. 4D) might increase MSH3 activity or increase the propensity of MSH3-mediated mismatch repair to generate repeat expansions, with a consistent effect across tissues and repeats as observed for coding variants that affect FAN1 function.

We also compared the modifiers of *TCF4* repeat expansion in blood to loci that influence risk of Fuchs endothelial corneal dystrophy (FECD), a common age-associated eye disorder that is thought to be caused (in most cases) by expansion of the *TCF4* DNA repeat^59,60^. Surprisingly, none of the loci we found to influence expansion of *TCF4* repeats in blood overlapped with loci that influence risk of FECD^61^, and none of our lead variants for *TCF4* instability (Table 1) associated with FECD (p>0.15) in a recent well-powered GWAS^61,62^. Additionally, FECD risk conferred by long *TCF4* repeats appeared to plateau for allele lengths beyond ∼75 repeat units (Supplementary Fig. 11). We did observe a modest effect of female sex on susceptibility for *TCF4* expansion in blood (∼10% higher odds of observing either a highly expanded allele or somatic expansion of a short allele; Fig. 4E). Further work will be required to determine whether this effect contributes to the >2-fold higher prevalence of FECD in females compared to males^63^, whether the instability-modifying genetic effects we identified are specific to blood (which is conceivable given the very different (more extreme) dynamics of *TCF4* somatic expansion in corneal endothelium^56^), and whether any modifiers of somatic expansion influence age at FECD onset.

### Repeat expansions in *GLS* associate with kidney and liver diseases

The deep phenotyping of the UKB cohort offered the opportunity to search for effects of repeat expansions on a wide variety of diseases and other clinical phenotypes. Expansions at four of the 15 repeat loci we studied (*GLS*, *HTT*, *TCF4*, and *DMPK*) associated with various diseases and biomarker measurements (Supplementary Tables 6 and 7). The associations of expansions in *HTT*, *TCF4*, and *DMPK* with disease phenotypes reflected the known roles of these repeats in HD, FECD, and myotonic dystrophy (DM1)^11^. However, associations of repeat expansions in *GLS* with biomarkers of kidney and liver function were surprising, as such expansions in *GLS* (which encodes kidney-type glutaminase) have only been observed to be pathogenic in extremely rare cases of severe, childhood-onset recessive glutaminase deficiency^64,65^. The large UKB population sample made it possible to see that heterozygous carriers of long *GLS* alleles exhibited anomalous phenotypes.

In UKB, 139 individuals (0.03%) carried a long CAG repeat (245 repeat units) within the 5’ UTR of *GLS* (Fig. 5A). Most of these individuals (98 of 139) exhibited evidence of a highly expanded allele (∼100+ repeats based on observing IRR pairs; Fig. 1B), consistent with long *GLS* repeats being highly unstable somatically^64^ and in the germline (Fig. 2B,D,E). Germline instability of the repeat increased rapidly with allele length: mid-length alleles (25–40 repeats) were already sufficiently unstable for mutations to be observable between close relatives, including a quartet of genetically-inferred first cousins among whom multiple intergenerational mutations had occurred (Fig. 5B).

**Figure 5:**
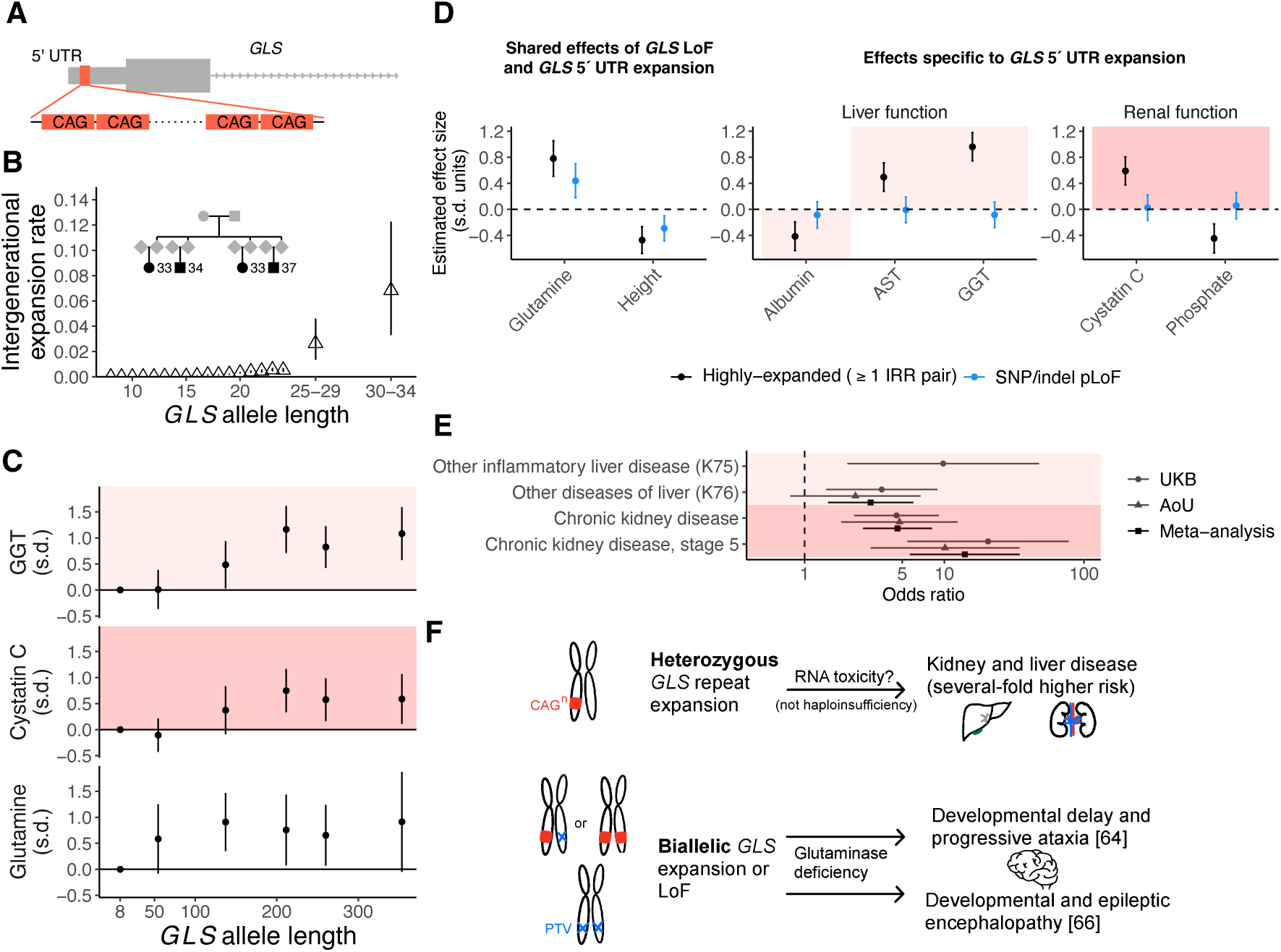
Instability and pathogenicity of CAG repeat expansions in the 5’ UTR of *GLS*. **(A)** Location of a polymorphic CAG repeat within the 5’ UTR of *GLS*. **(B)** Intergenerational expansion rates of short and mid-length *GLS* alleles estimated from IBD (for common alleles containing ≤24 repeat units) and from related pairs of UKB participants carrying rarer, mid-length alleles (25–34 repeat units). Inset, multiple mutations observed among four related UKB participants (likely to be mutual first cousins) carrying mid-length *GLS* repeat alleles. **(C)** Mean gamma-glutamyl transferase (GGT), cystatin C, and glutamine levels (adjusted for age, age squared, and sex) among individuals with long *GLS* alleles (stratified into quintiles) and individuals without a long allele (plotted at the modal length of 8 repeats). **(D)** Effect sizes of *GLS* alleles for liver and renal biomarkers and other quantitative traits measured in UKB, adjusted for age, age squared, and sex. *GLS* repeat expansions and SNP and indel variants predicted to cause loss of function (pLoF) all associated with increased glutamine levels (inverse-normal transformed) and decreased height; in contrast, only highly expanded *GLS* repeat alleles (≥1 IRR pair, i.e., ∼100+ repeat units) associated with altered serum biomarker levels. Shaded rectangles indicate biomarker effect directions typically associated with disease. **(E)** Increased odds of liver and renal diseases among UKB and AoU participants with highly expanded *GLS* repeat alleles compared to participants without a long allele. The stage 5 chronic kidney disease phenotype included ICD-10 codes for end-stage renal disease. Analyses in UKB adjusted for age and sex; analyses in AoU adjusted for, age, age squared, sex, and genetic ancestry. Error bars, 95% CIs. **(F)** Contrasting pathogenic effects and hypothesized mechanism for heterozygous *GLS* repeat expansions compared to biallelic *GLS* expansion or LoF.

Repeat expansions in *GLS* associated strongly with elevated biomarkers of liver and kidney disease (p=6.8×10^-15^ for gamma-glutamyl transferase (GGT); p=6.9×10^-7^ for cystatin C; Fig. 5C,D and Supplementary Table 6). These associations appeared to be driven by highly expanded alleles (beyond a threshold of ∼100–200 repeat units; Fig. 5C), and SNP and indel variants in *GLS* predicted to cause loss of function (pLoF) did not associate with these biomarkers (Fig. 5D and Supplementary Table 8). Carriers of highly expanded alleles exhibited several-fold increased risk of liver and kidney diseases, results that replicated in AoU, with particularly elevated risk of stage 5 chronic kidney disease (OR=14.0 [5.7–34.3], p=7.2×10^−9^ in meta-analysis across UKB and AoU; Fig. 5E and Supplementary Table 9). These results suggest that highly expanded *GLS* repeats cause a dominant (but low-penetrance) DNA-repeat disorder distinct from recessive glutaminase deficiency (Fig. 5F). Glutaminase deficiency is caused by biallelic impairment of *GLS* function—either by SNP/indel pLoF variants^66^ or long *GLS* repeats that suppress *GLS* expression^64^, both of which associated with elevated serum glutamine in UKB (Fig. 5C,D and Supplementary Table 8). In contrast, the effects on kidney and liver biomarkers were specific to highly expanded repeats, indicating a different pathological mechanism unrelated to GLS function, such as RNA toxicity^67^ (Fig. 5F).

## Discussion

These results show how biobank WGS data sets contain abundant information about DNA-repeat instability and its biological effects that can be accessed using new computational approaches and ideas. By developing a new approach to recognize somatic expansion from abundant WGS data, we identified new genetic modifiers of DNA-repeat expansion and observed repeat-specific effects of instability-modifying haplotypes at DNA repair genes.

Our results point to important tissue-specific differences in somatic expansion. While the strongest genetic associations with *TCF4* repeat expansion in blood involved several of the same genes that have consistently been identified by GWAS of other repeat-expansion-related phenotypes (e.g., age-at-onset of HD^18–22^, age-at-onset of X-linked dystonia-parkinsonism^53^, and *HTT* repeat expansion in blood^22^), the effect directions of associated haplotypes often differed, and the specific alleles responsible for these effects appeared to at least partially differ. These results reinforce recent evidence suggesting substantial tissue specificity of genetic modifiers of *HTT* expansion in blood versus brain^22^. Somatic expansion in brain appears to arise predominantly in specific types of neurons^24^ and thus involve non-replicative mechanisms, whereas long-term somatic expansion in blood is likely to arise from events in hematopoietic stem cells and thus may substantially arise in the course of DNA replication.

The clear and strong differences in genetic effects on repeat expansion in different tissues suggest a need for care and caution in efforts to use DNA repeats in clinically accessible tissues (such as blood) to inform on the status of somatic expansion in disease-relevant tissues (such as brain): to the extent that somatic expansion in blood is shaped by different alleles (even if at the same genes), then somatic expansion in blood may offer little information about somatic expansion in the brain beyond that both tend to increase with age, though it might still be a useful biomarker for the immediate effect of future therapies that seek to slow repeat expansion.

We also found that genetic modifiers of repeat instability can even act differently on expansions of different repeats in the same tissue (*TCF4* versus *HTT* expansion in blood). The modulation of genetic effects by locus-specific effects may suggest roles for locus-specific chromatinization or transcriptional dynamics and will be an interesting area for mechanistic studies.

The deep phenotype data available in biobank data sets also enabled us to discover a dominant DNA-repeat disorder involving highly expanded 5’ UTR repeat alleles in *GLS*, which associated with several-fold higher risk of kidney and liver diseases. Large WGS cohorts provide an opportunity to identify such pathogenic rare alleles that, despite their strong effects on disease risk, have not been discovered to date owing to their low penetrance in families. Analyses of the phenotypic effects of common repeat variation, which we did not undertake here, may uncover subclinical phenotypes and may also resolve the question of whether intermediate-length alleles of pathogenic repeats have any beneficial effects (that could in principle cause them to persist in human populations); association analyses conducted to date^9,10^ have not detected evidence of such effects.

Analysis of repeat instability in population biobanks does have several limitations. While we could study germline mutation rates by analyzing IBD among unrelated individuals, we could not assess effects of genetic variation, parental age, or parent-of-origin on germline mutability, as this requires ascertaining de novo mutations^14,17,33^. Additionally, the available short-read WGS data we analyzed provided only indirect glimpses of somatic mutation, through observations of solitary reads spanning mutated alleles, and through read-pair-based evidence of highly expanded alleles of unknown lengths. Nonetheless, the analytical tools we have developed here for biobank-scale WGS analysis provide a useful complement to studying repeat instability in families^13,14,17,33^ and in patient cohorts using targeted sequencing techniques^22,68^, and combining these approaches should provide opportunities for further discovery.

## Acknowledgments

We thank R. Cai and S. Browning for providing estimates of effective population size in UK Biobank, and we thank B. Gorman and S. Iyengar for helpful discussions. This research was conducted using the UKB resource under application no. 40709. M.L.A.H. was supported by a US National Institutes of Health (NIH) fellowship F32 HL160061. R.E.H. and S.A.M. were supported by US NIH grant R01 HG006855. N.K. was supported by US NIH training grant T32 HG002295 and fellowship F31 DE034283. R.E.M. was supported by US NIH grant K25 HL150334. S.R. was supported by a Swiss National Science Foundation Postdoc.Mobility fellowship. P.F.P was supported by ERC Starting Grant no. 850869. P.-R.L. was supported by US NIH grants R56 HG012698 and R01 HG013110 and a Burroughs Wellcome Fund Career Award at the Scientific Interfaces. The funders had no role in study design, data collection and analysis, decision to publish or preparation of the manuscript. The *All of Us* Research Program is supported by the NIH, Office of the Director: Regional Medical Centers: 1 OT2 OD026549; 1 OT2 OD026554; 1 OT2 OD026557; 1 OT2 OD026556; 1 OT2 OD026550; 1 OT2 OD026552; 1 OT2 OD026553; 1 OT2 OD026548; 1 OT2 OD026551; 1 OT2 OD026555; IAA no. AOD 16037; Federally Qualified Health Centers: HHSN 263201600085U; Data and Research Center:5 U2C OD023196; Biobank: 1 U24 OD023121; The Participant Center: U24 OD023176; Participant Technology Systems Center: 1 U24 OD023163; Communications and Engagement: 3 OT2 OD023205; 3 OT2 OD023206; and Community Partners: 1 OT2 OD025277; 3 OT2 OD025315; 1 OT2 OD025337; and 1 OT2 OD025276. In addition, the *All of Us* Research Program would not be possible without the partnership of its participants.

