## Supplementary material for "Insights into the causes and consequences of DNA repeat expansions from 700,000 biobank participants": Methods and supplementary material

### Methods and supplementary material for “Insights into the causes and consequences of DNA repeat expansions from 700,000 biobank participants”

Margaux LA Hujoel, Robert E Handsaker, Nolan Kamitaki, Ronen E Mukamel,
Simone Rubinacci, Pier F Palamara, Steven A McCarroll, Po-Ru Loh

**5 Supplementary Figures** **27**

**6 Supplementary Tables** **38**

**References** **47**

### 1 Expanded CAG repeats in UK Biobank

#### 1.1 Extracting WGS reads derived from long CAG repeats

To characterize the landscape of CAG repeat expansions in a generally-healthy population cohort, we searched for evidence of long CAG repeats in short-read, 151bp paired-end WGS data available for 490,416 UK Biobank participants. Specifically, we searched for in-repeat reads (IRRs), which we operationally defined as read sequences containing $\geq$45 distinct occurrences of a CAG repeat unit (i.e., CAG, AGC, GCA, CTG, TGC, or GCT). We chose the threshold of $\geq$45 to allow for base-calling errors while still ensuring that IRRs were nearly, if not completely, comprised of CAG repeats.

To efficiently identify such IRRs, we made use of the observation that conveniently, bwa [1] had aligned most IRRs (regardless of their locus of origin) to a single locus: the CAG repeat in *TCF4*, which is the longest CAG repeat in the GRCh38 reference genome (Supplementary Fig. 1A). More precisely, among 500 randomly selected WGS samples, 91% of all IRRs were aligned to *TCF4*, and among the remaining IRRs, nearly all either originated from *CA10* or contained a moderate amount of non-CAG flanking sequence and were therefore less relevant to our analyses (Supplementary Fig. 1B,C). This read-mapping behavior, which allowed us to extract most IRRs by analyzing only a single slice of each indexed WGS alignment (cram) file (reducing computational cost by $\sim$2,000-fold compared to scanning the entire cram file), was specific to alignments generated using bwa; in alignments generated by DRAGEN [2], IRRs were more frequently (correctly) mapped to the same loci as their mates.

Specifically, for each WGS cram file containing reads aligned to GRCh38 using bwa, we extracted reads aligned to chr18:55585150-55587230 (which contains the *TCF4* CAG repeat and 1kb of flanking sequence on either side). We excluded reads with any of the last four SAM flags set (`0xF00`). Among the subset of reads that aligned to the *TCF4* CAG repeat sequence (chr18:55586150-55586230), we analyzed read sequences to identify IRRs (as defined above). For each IRR, we further analyzed its mate to determine what information the paired reads provided about their originating sequence:

- If the mate aligned to the $\sim$2kb region chr18:55585150-55587230 (such that it had already been extracted):
  - If the mate was also an IRR, then we recorded the read pair as an "IRR pair" (indicative of a highly expanded repeat; Supplementary Fig. 1).
  - Else, if the mate had mapping quality (MAPQ) $>$ 30, then we recorded the read pair as an "anchored IRR" (Supplementary Fig. 1A) originating from the *TCF4* repeat.
- Else, if the mate aligned to a locus other than *TCF4* (i.e., to a different chromosome, before chr18:55580000, or after chr18:55590000), we extracted the mate from the cram file and

examined its mapping quality:

  - If the mate had MAPQ $> 30$, then we recorded the read pair as an anchored IRR (known to originate approximately from the mate's aligned position).

#### 1.2 Assigning in-repeat reads to CAG repeat loci of origin

Anchored IRRs provide only approximate information about locations of long repeats, as although one member of the read pair is aligned with high mapping quality, the exact distance between this location and the repeat itself is unknown (due to variability in WGS fragment lengths, which are typically several hundred base pairs but occasionally exceed 1kb). To identify the specific CAG repeat loci from which IRRs originated, we applied the following procedure:

1. Assign anchored IRRs to 100kb bins (based on Mb coordinates of aligned mates, rounded to one decimal place). Most 100kb bins did not contain an anchored IRR in any UKB participant; only 2844, 169, 38, and 27 bins contained an anchored IRR in at least 1, 2, 3, and 5 UKB participants, respectively.
2. Refine 100kb bins containing an anchored IRR in $\geq$5 UKB participants to specific CAG repeats previously genotyped in a high-quality STR reference panel [3]. We downloaded both per-locus summary statistics (including coordinates and repeat unit sequence) and population-specific statistics (including allele frequency and heterozygosity) from `https://github.com/gymrek-lab/EnsembleTR`. We then identified the subset of autosomal repeats with a CAG/CTG repeat unit that were polymorphic within the 1000 Genomes EUR population [4], comprising a total of 1,159 autosomal CAG repeats. Finally, for each of the 25 autosomal 100kb bins with an anchored IRR in $\geq$5 UKB participants, we computed the median (across participants) of the mean anchor location per participant, and we checked whether this median position was within 500bp of the start coordinate of a EUR-polymorphic CAG repeat. This procedure assigned 18 of the 100kb bins to specific autosomal CAG repeats (Supplementary Table 1).
3. Drop anchored IRRs with alignment locations not matching identified CAG repeats. For each of the 18 100kb bins that resolved to a CAG repeat in the previous step, we re-examined individuals with anchored IRRs in these 100kb bins to verify that their alignment locations were near the identified CAG repeat. This was nearly always the case; we found only three individuals with discrepant alignment locations (mean anchoring alignment position $>$10kb from the CAG repeat location) and dropped these individuals from the corresponding lists of anchored IRR carriers (one individual each for *CA10*, *LRMDA*, and *ATN1*).

We further assigned IRR pairs to CAG repeat loci from which they were likely to originate. Because these IRR pairs consisted completely or nearly completely of CAG repeat sequence, they could not be confidently aligned to loci of origin based on their sequence composition. However, any locus containing a highly expanded repeat generating an IRR pair would be expected to also generate several anchored IRRs (Supplementary Fig. 1). Because most individuals had at most one locus generating many anchored IRRs, we could therefore assign IRR pairs to loci of origin with high confidence by examining counts of anchored IRRs aligned to 100kb bins.

Specifically, if all of an individual's anchored IRRs aligned to the same 100kb bin, we assigned that individual's IRR pairs to this bin. If an individual had anchored IRRs aligned to multiple bins, we assigned IRR pairs to the bin containing the largest number of anchors, excluding the *CA10* locus (because although long *CA10* alleles are common, highly expanded *CA10* alleles are rare: among individuals with anchors only at *CA10*, only ∼1% (1,069 of 122,238) had IRR pairs). In the rare case of a tie between multiple 100kb bins (other than *CA10*) having the same largest number of anchors, we randomly assigned IRR pairs to tied bins.

##### 1.3 Quantifying enrichment of repeat expansions in transcribed regions

Most of the 18 CAG repeat loci for which we detected long alleles in ≥5 UKB participants were in exons. To test whether repeat expansions were enriched in various transcriptional contexts, we intersected the list of 1,159 EUR-polymorphic autosomal CAG repeats with GENCODE v44 canonical transcript annotations to determine the transcriptional context of repeats. We then used Fisher's exact test to assess whether CAG repeats in a given context were more or less likely to be expanded to ≥45 repeat units in ≥5 UKB participants (Supplementary Table 2).

##### 1.4 Selection of CAG repeat loci for downstream analysis

Among the 18 CAG repeat loci identified by the above analyses (Supplementary Table 1), we took forward a subset of 15 CAG repeats for further analysis of germline and somatic mutability based on satisfying the following criteria: (i) heterozygosity >0.01 in EUR [3], (ii) >5 individuals with anchored IRRs in the repeat's 100kb bin, and (iii) only one CAG repeat segment contained within the repeat structure (for ease of genotyping).

#### 2 Intergenerational instability of CAG repeats

##### 2.1 Identifying short and mid-length alleles from spanning reads

For each of the 15 selected CAG repeat loci, we identified *short and mid-length* alleles (<45 repeat units) supported by spanning reads. We used the following pipeline to identify alleles observed within each UKB participant, recording both alele lengths and intra-repeat sequence variations:

1. Extract reads aligned to the repeat locus with mapping quality $\geq$30 and with mate mapped to the same chromosome.
2. Restrict to spanning reads containing exactly one occurrence each of a starting and ending flank sequence (9bp, consisting of one 3bp repeat unit along with 6 flanking base pairs directly before and after the repeat in the GRCh38 reference genome; Supplementary Table 1).
3. For each allele length supported by a spanning read, determine a consensus sequence for this allele (in the individual being analyzed). Specifically, assign each base of the consensus repeat sequence to be the most commonly-observed base at that position among all spanning reads supporting the repeat length under consideration, restricting to base calls with quality $\geq$25 (i.e., QUAL of `F` or `:` under the base-quality discretization used in UKB cram files). Set bases with no high-quality base calls or with a tie for the most common base to missing.

To generate the allele frequency histograms shown in Fig. 1B, we computed allele frequencies among individuals we could confidently genotype as heterozygotes (based on observing two distinct alleles with $\geq$5 spanning reads and no other alleles with $>$2 spanning reads) or were likely to be homozygotes (based on observing a single allele with $\geq$10 spanning reads and no other alleles with $>$2 spanning reads).

#### 2.2 Estimating germline mutation rates of short alleles

We estimated repeat-specific, allele-specific intergenerational expansion and contraction rates by analyzing length discordances among alleles belonging to genomic tracts inherited identical-by-descent (IBD), building upon IBD-based analyses of single-nucleotide mutations [5–9]. We restricted these analyses to common, shorter alleles ($\leq$30 repeat units) that were typically spanned by several short reads, such that alleles were unlikely to "drop out" due to not generating any spanning reads by chance.

For each of the 15 CAG repeat loci, we performed analyses within a (locus-specific) subset of individuals heterozygous for short alleles that could be phased with high confidence. That is, we required that individuals have:

- Two distinct alleles each supported by $\geq$5 spanning reads and no other alleles with $>$2 spanning reads.
- Length difference of $\geq$3 repeat units separating the two alleles (i.e., $|A_1 - A_2| \geq 3$), facilitating phasing of the alleles onto the individual's two haplotypes.

We further restricted analyses to individuals with SNP-array genotypes [10] for whom we had previously generated phased haplotypes and identified longest IBD matches [11]. Among these individuals, we performed the following sequence of analyses:

1. **Phase repeat alleles onto SNP-haplotypes** (using IBD matches to determine which of an individual's two haplotypes contains the longer allele). We used an iterative algorithm starting by randomly assigning phase and then running 50 phase-update iterations: for each individual in turn, we computed the mean allele length across the longest five IBD matches for haplotype 1 (respectively, haplotype 2) and then re-assigned phase according to which haplotype was estimated to carry a longer allele.
2. **Identify recent IBD pairs and the ancestral allele of each pair.** For each haplotype of each individual, we examined the haplotype's longest IBD match. If the IBD length was $>$5 cM and spanned $>$0.5 cM on each flank of the CAG repeat locus, we attempted to identify the ancestral allele of the IBD pair using "outgroup" haplotypes sharing less-recent IBD (Fig. 2A):

    - For each of the two haplotypes in the IBD pair, identify its longest 20 IBD matches. Define the set of outgroup haplotypes to be the intersection of these two top-20 sets.
    - If $\geq$5 outgroup haplotypes are found, examine the repeat alleles carried by the top five outgroup haplotypes. If a majority of these alleles ($\geq$3 of 5) match one another (in allele length), and if this consensus allele also matches one or both of the alleles in the IBD pair, designate this consensus allele to be the ancestral allele of the IBD pair.

3. **Determine an approximately independent set of inferred allele transmissions** (from ancestral alleles to IBD pairs). For each IBD pair with an ancestral allele identified in the previous step, we computed the allele length difference ($\Delta$) between each present-day allele (in the IBD pair) and the ancestral allele. One or both of the alleles was required to have $\Delta$=0 in the previous step. We further required that $\Delta \in \{-2, -1, 0, 1, 2\}$ (reasoning that larger jump sizes might be more likely to arise from genotyping or phasing error). Finally, we removed any duplicate IBD pairs: we had ascertained IBD pairs by examining each haplotype's longest IBD match, such that closely-related haplotypes often nominated each other. This final set of IBD pairs and their inferred ancestral alleles represented an approximately non-redundant set of allele transmissions, as longest IBD matches should typically correspond to leaf-pairs on a coalescent tree.
4. **Estimate the number of generations between each ancestral allele and IBD pair.** To do so, we computed the expected time to the most recent common ancestor (TMRCA) of each IBD pair based on the length of the shared IBD tract and a demographic model accounting for recent population growth.

    If two alleles have a known TMRCA, the distribution of the length (in Morgans) of the IBD segment containing them is given by a gamma distribution with shape parameter 2 and rate parameter $2t$ per Morgan ($f(L|\mathrm{TMRCA} = t) = L(2t)^2e^{-2tL}$; pages 57–58 of [12]). To compute expected TMRCA given the length of an IBD tract, we can use Bayes' theorem to obtain $P(t|L) \propto f(L|t)P(\mathrm{TMRCA} = t)$. Thus, if $L$ is the length of an IBD segment in

Morgans and TMRCA $= t$,

$$\mathbb{E}[t|\text{IBD length} = L] = \frac{\int t^3 e^{-2tL} P(\text{TMRCA} = t)dt}{\int t^2 e^{-2tL} P(\text{TMRCA} = t)dt} \tag{1}$$

The term $P(\text{TMRCA} = t)$, which represents a prior on the TMRCA of two alleles, is dependent on demographic history. This probability distribution can be numerically computed assuming that the effective population size $N_e(t)$ at each generation $t$ in the past is known: under a coalescent model, coalescence occurs with probability $1/(2N_e(t))$ at generation $t$ (assuming it has not yet occurred), allowing iterative computation of $P(\text{TMRCA} = t)$ for $t = 1, 2, 3, \ldots$. We used autosomal $N_e(t)$ values that had previously been estimated for the past 300 generations based on an analysis of 10,000 White British UKB participants [13].

5. **Estimate allele-specific per-generation expansion and contraction rates.** For a given allele $A$, the estimated rate of intergenerational mutational jumps of size $c$ is $\frac{\sum(\Delta==c)}{\sum \text{TMRCA}}$, where the sums are across transmissions within IBD pairs with ancestral allele $A$. This calculation assumes that at most one mutation occurred between each present-day and ancestral allele, which is a reasonable assumption here (for IBD >5 cM and mutation rates <0.01 per generation).
6. **Estimate confidence intervals of mutation rates.** To do so, we rounded the total number of generations ($\sum \text{TMRCA}$) to the nearest integer and computed a binomial confidence interval.
7. **Recalibrate mutation rates and CIs to adjust for ascertainment of longest IBD matches.** The procedure above slightly underestimated mutation rates because the above computation of $\mathbb{E}[\text{TMRCA}|\text{IBD length} = L]$ did not account for our ascertainment of only the longest IBD match for each haplotype. (Intuitively, knowing that an IBD match of length $L$ was the longest IBD match for a given haplotype means that its expected TMRCA is more recent than that of a random IBD match of length $L$.) To correct for this issue, we computed an approximate calibration factor estimating the extent to which we had overestimated TMRCAs. To do so, we compared mutation rates computed using the above procedure to corresponding computations in which we relaxed the requirement that each IBD pair be the longest IBD match for a haplotype, instead considering all IBD pairs with length >5 cM spanning >0.5 cM on each flank of the CAG repeat locus. (This larger set of IBD pairs contains some redundancy of allele transmissions, making confidence intervals difficult to estimate, but removes ascertainment bias.) Across the 15 CAG repeat loci, we observed that we had underestimated mutation rates by a median factor of 1.204548 (computed by comparing mean rates of $\Delta = \pm 1$ mutations, aggregated across ancestral alleles). We therefore multiplied all mutation rate estimates and confidence interval sizes by this correction factor.

#### 2.3 Validating germline mutation rate estimates

We validated our approach to estimating germline mutation rates using two strategies. First, we examined the empirical probability of observing a length discordance between the two alleles in an IBD pair as a function of their expected TMRCA. If such discordances primarily reflect mutations (rather than genotyping errors or errors in determining IBD), and if expected TMRCA is correctly estimated, then the probability of discordance should scale approximately linearly with TMRCA (e.g., the probability of a mutation occurring among 20 allele transmissions should be roughly twice that of a mutation among 10 allele transmissions). In contrast, error modes that generate either false-positive discordances or noise in estimated TMRCA should flatten this relationship and produce a nonzero intercept at TMRCA=0. We implemented this analysis on the set of IBD pairs used in our germline mutation rate analyses and computed, for each decile of expected TMRCA (adjusted for ascertainment), the fraction of IBD pairs for exhibiting a length discordance of 1 repeat unit (as our analyses focused on 1-repeat-unit expansions and contractions). Across all 15 CAG repeat loci, we observed that discordance probabilities did indeed scale approximately linearly with TMRCA, as expected (Supplementary Fig. 2A).

Second, we directly validated germline mutation rate estimates using IBD2 siblings (i.e., siblings who inherited the same maternal haplotype and the same paternal haplotype at a given locus). The UKB cohort contained ~5,000 IBD2 sib pairs per locus, and while this sample size was insufficient to validate allele-specific mutation rates, it was large enough to provide a check on the population-average mutation rate of each CAG repeat locus (i.e., the probability that a randomly-sampled allele mutates in one generation). We first computed this value using all IBD pairs used in our germline mutation rate analyses (by dividing the total number of 1-repeat-unit discordances by twice the total TMRCA, adjusted for ascertainment). We then computed the corresponding value within IBD2 sibs, applying an analogous set of filters to the individuals included in analysis (i.e., restricting to confidently-phased heterozygotes and restricting to allele pairs that differed in length by $\Delta \in \{-2, -1, 0, 1, 2\}$) and then computing an analogous population-average mutation rate (by dividing the total number of 1-repeat-unit discordances by twice the number of allele pairs, since each allele pair had TMRCA=1). We computed a confidence interval for the IBD2-sib-based estimate using an exact binomial test as implemented in `binom.test` in `R` [14] and observed that for all 15 CAG repeat loci, the IBD2-sib-based confidence interval contained the estimate based on our full analysis, and the point estimates from the two approaches were broadly consistent (Supplementary Fig. 2B).

#### 2.4 Estimating germline mutation rates of mid-length *GLS* alleles

Common, short alleles of the *GLS* 5' UTR repeat (≤24 repeat units) exhibited particularly high germline mutation rates relative to the other CAG repeats we analyzed, leading us to wonder how much more quickly rare, mid-length *GLS* alleles (25–40 repeat units) might mutate. Assessing the mutation rates of these longer alleles using the analysis pipeline described above was not feasible because of the rarity of these alleles and the smaller numbers of spanning reads typically available for genotyping them (due to their longer lengths). Instead, we studied the mutation rates of these alleles using a separate pipeline, analyzing allele length discordances among close relatives in UKB based on DRAGEN genotyping [2] (which more optimally utilized available read data).

We first identified IBD pairs with mid-length alleles using the following procedure. For each individual that DRAGEN had genotyped to carry one allele of length $\geq$25 repeat units (supported by $\geq$1 spanning read) and another allele of length $<$20 repeat units, we identified the five longest IBD matches of each of the individual's two haplotypes. We phased the individual's short allele onto the haplotype for which the larger fraction of individuals sharing longest IBD with that haplotype carried the short allele. The mid-length allele was thus phased onto the opposite haplotype, such that the top IBD matches of that haplotype constituted IBD pairs of interest. For each such IBD pair, we assumed that the other individual's longer allele was the shared allele (as the large majority of *GLS* alleles are short; mid-length and long *GLS* alleles are rare).

We then analyzed the subset of these IBD pairs that involved close relatives (third-degree or closer based on previously-computed kinship coefficients $\phi$ [10]). To estimate germline mutation rates, we divided the number of discordances among IBD pairs by the sum of the numbers of generations separating these pairs, estimated as $-\log_2(\phi)$ (reasoning that full-sib, avuncular, and first-cousin relationships were likely to comprise most such relationships). We estimated germline mutation rates separately for haplotype-pairs for whom the shortest mid-length allele among the pair had length 25–29 repeats or 30–34 repeats. As above, we estimated confidence intervals by rounding the total number of generations to the nearest integer and computing a binomial confidence interval, which was a reasonable approximation given that only a small fraction of individuals appeared in multiple pairs of closely related haplotypes.

### 3 Somatic instability of CAG repeats

#### 3.1 Quantifying somatic instability of short CAG repeat alleles

An initial analysis of common, short alleles of the *GLS* repeat ($\leq$24 repeat units) also showed clear evidence of cases of somatic mosaicism: for 15 UKB participants, WGS data indicated the presence of three distinct alleles (each supported by $\geq$5 spanning reads), and these individuals were $\sim$5 years older than average (mean age of 61.3 years). This observation led us to try to quantify somatic instability of alleles short enough ($\leq$30 repeat units) to typically be spanned by several WGS reads.

##### 3.1.1 Filtering aberrant spanning reads arising from PCR stutter errors

We hypothesized that we might be able to find evidence of somatic mutation from single spanning reads suggesting a $\pm1$ or $\pm2$ repeat unit mutation relative to an inherited allele (supported by many spanning reads). The main challenge of such an analysis is that such aberrant reads commonly arise from "PCR stutter" error caused by polymerase slippage during PCR amplification [15–17]. This error mode arises even in WGS data generated from libraries prepared using PCR-free protocols (which were used by UKB and AoU) because of the "bridge amplification" step of sequencing by synthesis, in which a DNA fragment to be sequenced is PCR-amplified into a localized cluster of (usually identical) single-stranded DNA molecules anchored to a flow cell [18] (Fig. 2C). Sequencing is then performed by iteratively adding fluorescently tagged nucleotides complementary to these single-stranded DNA molecules (beginning with a primer and extending the complementary DNA strand one base pair at a time) and measuring the fluorescence signal generated (in aggregate) by the nascent nucleotide of each molecule in the cluster. At each successive iteration, a base call (corresponding to the nucleotide position just added) is derived from this fluorescence signal.

When bridge amplification occurs without error, the DNA molecules that form each clonal cluster are identical, such that at each iteration of sequencing by synthesis, the same new nucleotide is added to the complementary DNA strand of each molecule in the cluster, and the same fluorescence signal is emitted from each newly-added nucleotide in the cluster, generating a high-quality base call. However, if a PCR stutter error occurs during bridge amplification, a polyclonal cluster containing a mixture of molecules (some containing the error and others not containing the error) is produced (Fig. 2C). Consequently, at some base positions, molecules with and without the error will contain different nucleotides, such that the aggregate fluorescence signal generated by the polyclonal cluster will contain a mixture of signals corresponding to the distinct nucleotides represented in the cluster. In this scenario, the base call that is made might correspond to either the original sequence or the error-modified sequence, and—importantly for our purposes—the base quality (i.e., base call confidence) will be reduced.

The upshot of the above behavior is that when an aberrant spanning read (supporting a repeat allele that appears to harbor a length mutation) is generated by a PCR stutter error during bridge amplification, the base quality string corresponding to this read sequence should contain a *predictable* sequence of low-quality bases: read positions through the end of the repeat should have high quality (because the two species of DNA molecules present agree up to this point), after which positions at which the molecules with and without the error have inconsistent nucleotides should have low quality (Fig. 2C). Examining base qualities of aberrant spanning reads (some of which reflect real somatic mutations, but many of which arise from PCR stutter errors) showed that for many reads, this pattern was indeed readily visible (Fig. 2C and Supplementary Fig. 4A).

Based on these observations, we devised a filtering strategy to identify a high-quality set of aberrant spanning reads in which real somatic mutations were enriched and PCR stutter errors were depleted. The main idea of this filtering strategy was to identify reads with high base qualities at the positions at which PCR stutter error would be expected to reduce base qualities. This filter was not expected to be perfect, particularly because polymerase slippage during the initial step of bridge amplification (in which the original DNA fragment to be sequenced hybridizes to an oligo on the flow cell, a polymerase creates the complement of the hybridized fragment, and the original fragment washes away) produces a monoclonal cluster in which all DNA molecules contain the PCR stutter error, leaving no evidence of error in the base quality string. Nonetheless, for repeats with relatively higher somatic mutation rates, and for alleles of longer lengths (which are more prone to somatic mutation), we reasoned that somatic mutations could be sufficiently common to represent the majority of aberrant reads that passed stringent filtering, allowing analysis of somatic mutation of short repeat alleles from UKB and AoU WGS data.

Specifically, we applied the following analysis pipeline to each CAG repeat locus:

- Restrict to heterozygous individuals with exactly two alleles supported by $\geq$3 spanning reads, with these two alleles differing in length by $\geq$5 repeat units ($|A_1 - A_2| \geq 5$). In such individuals, we could confidently determine which allele a putative mutation (supported by an aberrant spanning read) arose from.
- Identify aberrant alleles differing in length from an inherited allele by $\pm1$ or $\pm2$ repeat units and supported by exactly one aberrant spanning read.
- Filter aberrant spanning reads to those with high base qualities at diagnostic positions and with robustly-determined allele lengths. For each aberrant spanning read, we assumed that the aberrant allele originated (either due to somatic mutation or PCR stutter error) from the inherited allele closer in length. We then determined which diagnostic read positions would be expected to have low quality had the aberrant allele been produced during bridge amplification. To do so, we shifted the sequence of the appropriate flank—right flank for reads aligned in forward orientation; left flank for reads aligned in reverse orientation—by $\pm$3bp or $\pm$6bp toward the repeat (corresponding to the other sequence that would be expected to contribute to the cluster, had a PCR stutter error occurred) and identified positions at which the unshifted sequence differed from the shifted sequence (Fig. 2C). Finally, we required that the aberrant spanning read satisfy all of the following filters:
  - $\geq$4 diagnostic base positions identified.
  - Maximal base quality (`F`, i.e., Q score of 37) observed at $\geq$80% of diagnostic positions.
  - Perfect or near-perfect match to the expected sequence on the opposite flank (left flank for forward reads; right flank for reverse reads). This filter ensured that base positions sequenced before the repeat sequence were accurately sequenced and were consistent with the read truly being a spanning read. Specifically, we required $\geq$95% sequence identity (compared to the consensus sequence of the originating inherited allele) among

high-quality bases (Q≥25; i.e., QUAL of `F` or `:`) on the first-to-be-sequenced flank.

– High base quality (`F` or `:`) at any key bases at which a single miscalled base would result in incorrect allele sizing. For example, the left flank of the *TCF4* repeat contains the sequence AGGAG**G**AG**C**AGC; the two bolded bases are key bases because a G-to-C base-calling error in the first base would result in an additional AGC repeat unit, and a C-to-G error in the second base would result in one fewer AGC repeat unit.

##### 3.1.2 Estimating fractions of blood cells harboring somatic mutations

The stringent filtering procedure described above identified a set of aberrant spanning reads free of detectable evidence of PCR stutter error and thus representing putative somatic mutations. However, to use these reads to quantify the propensities of different alleles to mutate somatically (e.g., the rate at which an allele of length $L$ repeats mutates somatically to an allele of length $L+1$), we needed to compute a normalization factor. Specifically, we needed to know how many reads we would expect to pass the same stringent filters in a heterozygous individual who inherited an $L$-repeat allele but in whom this allele had mutated to length $L+1$ in 100% of blood cells. Given this latter quantity, we could then estimate the fraction of blood cells in which an $L \to (L+1)$ mutation had occurred (averaged across biobank participants who inherited a single $L$-repeat allele) by dividing the mean number of aberrant spanning reads supporting an $L \to (L+1)$ mutation (and passing filtration) by this denominator.

To estimate the above denominator, we identified individuals who were heterozygous for a presumably-inherited allele of length $L+1$ and then filtered spanning reads supporting the $(L+1)$-repeat allele as if we were evaluating them for having arisen from somatic mutation or PCR stutter error modifying an $L$-repeat allele. The average number of $(L+1)$-repeat-spanning reads that passed filtration (per heterozygous carrier of an $(L+1)$-repeat allele) gave an estimate of the desired denominator.

Putting this all together, we estimated the fraction of blood cells harboring an $L \to (L+1)$ mutation as:

$$\frac{\text{Mean number of aberrant reads supporting an } L \to (L+1) \text{ mutation and passing filtration}}{\text{Mean number of reads spanning an } (L+1)\text{-repeat inherited allele and passing filtration}} \tag{2}$$

where the mean in the numerator is across the subset of individuals heterozygous for an $L$-repeat allele described above, and the mean in the denominator is across individuals heterozygous for an $(L+1)$-repeat allele.

The same approach was also applicable for estimating fractions of blood cells in which $L$-repeat alleles had mutated to alleles of length $L-1$ or $L \pm 2$, but only somatic mutations to length $L+1$ appeared to occur sufficiently frequently relative to PCR error to obtain robust results.

##### 3.1.3 Identifying CAG repeat loci with evidence of somatic expansion

To assess which of the 15 CAG repeat loci we analyzed showed evidence of somatic instability of common, shorter alleles, we examined the fraction of cells (among UKB participants heterozygous for a given allele) estimated to harbor a 1-unit expansion (respectively, 1-unit contraction) of each common allele of length $\leq$30 repeats (Supplementary Fig. 5). For most CAG repeat loci, estimated fractions of cells with putative contractions exceeded those with putative expansions, suggesting that residual PCR stutter error (which is contraction-biased [15]) could be dominating these estimates. However, for four loci (*GLS*, *ATN1*, *TCF4*, and *DMPK*), putative expansions began to outnumber putative contractions for less-short alleles, typically beyond an allele length of $\sim$15 repeats.

To formally test each CAG repeat locus for evidence of somatic expansion (respectively, contraction), we tested whether the age of a UKB participant associated with whether or not an individual's WGS data contained an aberrant spanning read that passed filtration and supported a 1-repeat-unit expansion (respectively, contraction). We restricted this analysis to UKB participants with at least one allele of length $>$15 and included the length of each individual's longer allele as a covariate.

Three loci exhibited Bonferroni-significant associations ($p < 0.05/15$) between age and the presence of a sequencing read putatively derived from a 1-repeat-unit somatic expansion (*GLS*, *TCF4*, and *DMPK*). Additionally, we noticed that *ATN1* apppeared to exhibit a strong increase in estimated somatic expansion rate and an expansion-vs.-contraction skew at slightly longer allele lengths (Supplementary Fig. 5), so we looked more carefully for an age effect in the precise allele range of potential somatic expansion ($\geq$18 repeats) and observed a significant association with age ($p < 0.05$).

##### 3.1.4 Assessing efficacy of stringent filtering to deplete PCR stutter errors

We performed two analyses to assess the effect our filtering approach. First, for each allele length $L$ at each CAG repeat locus, we evaluated the impact of the filter on our estimate of the fraction of blood cells harboring an $L \to (L+1)$ mutation. That is, we compared the estimate we obtained using the analytical pipeline described above to an analogous computation in which we did not attempt to filter PCR errors (i.e., counting all spanning reads supporting an $(L+1)$-repeat allele in both the numerator and denominator of equation (2)). As expected, filtering reduced estimated rates of somatic mutation (Supplementary Fig. 4B), indicating that PCR stutter errors were a source of aberrant spanning reads.

Second, we evaluated the efficacy of the filter in enriching for true somatic mutations among aberrant spanning reads. As noted above, we did not expect the filter to completely eliminate PCR stutter error (particularly because errors introduced during the initial step of bridge amplification are undetectable), so we wished to estimate what fraction of aberrant spanning reads that survived filtering represented real somatic mutations (rather than technical artifacts).

To do so, we analyzed the extent to which the estimated fraction of blood cells harboring an $L \rightarrow (L+1)$ mutation increased with age. Assuming that such somatic mutations accrue at an approximately constant rate with age, we would expect the fraction of cells carrying a mutation to scale linearly with age, whereas technical artifacts that produce aberrant spanning reads should be observed at a frequency independent of age. We therefore regressed $y =$ estimated fraction of blood cells harboring an $L \rightarrow (L+1)$ mutation on $x =$ age, including an intercept term in the regression, and we then divided the intercept by the mean $y$-value to obtain the relative contribution of technical artifacts to the $y$-values we had measured. Subtracting this quantity from 1 gave an estimate of the effectiveness of our filtering strategy.

To reduce noise in these regressions, we restricted analyses to alleles with >1,000 carriers, and we performed analyses on groups of alleles of lengths $L \leq$10, 11–15, 16–20, 21–25, and ≥26 repeats. Additionally, we performed these analyses in the *All of Us* cohort as its age range (18–90+ years) was much wider than UKB (40–70 years), facilitating regression analysis on age. These analyses indicated that for *GLS* and *TCF4*, the majority of aberrant spanning reads that passed filtering were indeed of somatic origin, and the same was true for longer (more mutable) alleles of *ATN1* and *DMPK* (Supplementary Fig. 4C).

##### 3.1.5 Replicating age-associated somatic instability of CAG repeats in *All of Us*

To evaluate the robustness of our estimates of fractions of blood cells harboring 1-repeat-unit somatic expansions, we compared estimates we obtained by analyzing WGS data from the UKB and AoU cohorts. We subdivided each cohort into three age tranches and computed estimates within each age tranche for each of the four CAG repeat loci that we had found to exhibit detectable somatic instability in blood (*GLS*, *ATN1*, *TCF4*, and *DMPK*). We observed concordant estimates between the two cohorts, and in both cohorts, somatic expansions were estimated to be more frequent among individuals of older ages (Supplementary Fig. 6).

#### 3.2 Quantifying somatic instability of long CAG repeat alleles at *TCF4*

Some CAG repeat loci are known to be somatically unstable, such that observations of highly expanded alleles (∼100+ repeats, producing IRR pairs) might indicate somatic expansion. To see if we had evidence of somatic expansion in UKB, we computed the average age among carriers of highly expanded alleles of each CAG repeat. In UKB, carriers of highly expanded *TCF4* repeat alleles were significantly older than average (+2.48 years; s.e. 0.08 years), suggesting a contribution of somatic expansion to these particularly long alleles. *TCF4* was the only locus for which carriers of highly expanded alleles were significantly older than average, so we focused further analyses of somatic instability of long repeats on the *TCF4* locus.

##### 3.2.1 Estimating lengths of long *TCF4* repeats

To more directly assess somatic expansion of long *TCF4* repeat alleles in UKB and AoU, we needed a way to estimate lengths of these alleles from short-read WGS data. Counts of in-repeat reads (IRRs) provided a straightforward way to do so, as the number of such reads observed in an individual heterozygous for a long allele (≥45 repeat units) increases approximately linearly with the length of the allele and with autosomal sequencing coverage:

$$\text{Estimated allele length (in repeat units)} = 45 + (\text{\# IRRs}) \times \frac{X}{\text{coverage}} \times \frac{1}{3}, \tag{3}$$

where 45 is the number of repeat units required for an allele to produce IRRs (according to our definition of IRR), $\frac{X}{\text{coverage}}$ is a constant calibration factor estimating that a new read is generated every $\frac{X}{\text{coverage}}$ base pairs of *TCF4* repeat sequence, and $\frac{1}{3}$ converts base pairs to repeat units. For an individual who is mosaic for long alleles of different lengths, this formula estimates the mean length across alleles present in the sequenced DNA sample.

We estimated the calibration factor $X$ by comparing short-read and long-read sequencing data among heterozygous carriers of long *TCF4* alleles for whom long-read data were available in AoU v7. Specifically, we set the median repeat length estimated from long-read data to equal the median repeat length estimated from short-read data (using the above formula), which gave $X \approx 424$. We used the following procedure to measure *TCF4* repeat lengths from long reads. For each carrier of a long *TCF4* allele (identified from short-read analysis), we identified long reads mapping to the *TCF4* repeat in GRCh38 with mapping quality ≥30, excluding reads with any of the last four SAM flags set (`0xF00`). We extracted the segment of each read containing the CAG repeat sequence by searching for exact matches to the left and right flanks of the repeat sequence (AGGAGGAG**CAGCAG** and **CAGCAG**CATGAAA); we verified that this approach generally gave the same result as identifying the first and last occurrences of a string of five CAGs (CAGCAGCAGCAGCAG). We then estimated the length of the repeat allele by computing the number of base pairs between the left and right flanks, dividing by 3, and adding 1. To guard against the possibility of a technical artifact producing extraneous, non-CAG sequence between the flanks, we required that this estimate agree with the number of CAG substrings found within the read sequence to within 50 repeat units.

##### 3.2.2 Estimating relative lengths of inherited *TCF4* repeat alleles

The length of a long *TCF4* repeat allele (estimated from short-read WGS data as described above) reflects both the effect of somatic expansion and the length of the allele that was originally inherited. To increase statistical power to analyze somatic expansion of *TCF4* repeats, we wished to control for inter-individual variation in inherited *TCF4* allele lengths. Doing so directly was not possible given that DNA sequencing data was available from a single time point per individual. Instead, we approximately estimated the relative length of each of an individual's two inherited *TCF4* repeat alleles (relative to the alleles inherited by other sequenced individuals) by using statistical imputation from other individuals who shared an extended—typically multi-megabase—SNP-haplotype at *TCF4* (and were therefore likely to have co-inherited a *TCF4* repeat allele of the same or similar length). Measurements of *TCF4* repeat alleles in these distantly-related individuals were themselves inexact estimates of the lengths of the alleles that these individuals had inherited (both because of sampling noise and because of somatic expansion), but we reasoned that averaging across multiple reference individuals would mitigate the impact of noise, and that while imputed allele lengths overestimate the lengths of longer inherited alleles more so than less-long inherited alleles (due to greater rates of somatic expansion), they still provide a useful measure of the relative lengths of inherited alleles. In detail, the imputation pipeline proceeded as follows:

1. **Computing coverage-adjusted counts of in-repeat reads (IRR).** As noted above, the number of IRRs generated by a *TCF4* repeat allele increases approximately linearly with the length of the allele and with WGS coverage, such that coverage-adjusted IRR count provides a quantification of allele length. In our imputation pipeline, we computed coverage-adjusted IRR count using a slightly different method than described above (simply because we ran the imputation analysis before working out details of equation (3)):
   - In UKB, we computed each sample's coverage-adjusted IRR count (for *TCF4*) as $(\#_{\text{anchored IRRs at } TCF4} + 2 \times \#_{\text{IRR pairs}})/(\text{cov}_{\text{GC67}}/\overline{\text{cov}_{\text{GC67}}})$, where $\text{cov}_{\text{GC67}}$ denotes a WGS sample's mean autosomal coverage in 400bp windows with 67% GC content (following the GC normalization approach of Genome STRiP [19]), and $\overline{\text{cov}_{\text{GC67}}}$ denotes the mean of this value across UKB WGS samples. We set coverage-adjusted IRR count to 0 (overriding the above formula) if an individual either had no anchored IRRs at *TCF4* or had been genotyped to be heterozygous for two short alleles.
   - In AoU, we normalized using the genome-wide coverage metric provided with the AoU v7 WGS data release; i.e., we computed $(\#_{\text{anchored IRRs at } TCF4} + 2 \times \#_{\text{IRR pairs}})/(\text{cov}/\overline{\text{cov}})$ and set this value to 0 if an individual had no anchored IRRs at *TCF4*.

2. **Identifying and phasing carriers of long *TCF4* alleles.** We defined "long-allele carriers" as individuals with nonzero coverage-adjusted IRR count, and in each data set (UKB and AoU), we created an imputation reference panel comprised of the subset of long-allele carriers who had no anchored IRRs at any locus other than *TCF4* or *CA10* (such that IRR pairs could be assumed to nearly always be derived from *TCF4*). We defined SNP-haplotypes of these individuals in the genomic region surrounding *TCF4* using our previous phasing of the UKB

SNP-array data set [20] and using `phase_common` from SHAPEIT5 [21] to phase the AoU v7 SNP-array data set. We then determined which haplotype(s) of each long-allele carrier contained a long *TCF4* allele using the following phasing approach:

- For each of an individual's two haplotypes $i = 1, 2$, we identified the top 20 longest SNP-haplotype matches at *TCF4* in the reference panel (quantified using IBS length as in ref. [22]), and we then computed the fraction $f_i$ of long-allele carriers among the reference individuals in whom these top 20 SNP-haplotype matches were found.
- If both $f_1 > 0.5$ and $f_2 > 0.5$, we considered the individual to be homozygous-long.
- Otherwise, we considered the individual to be a heterozygous carrier of a single long allele, and we assigned the long allele to the haplotype $i$ with higher $f_i$.

3. **Computing an inherited *TCF4* allele length metric by imputing coverage-adjusted IRR count**. For each long-allele carrier, for each haplotype that we determined above to carry a long *TCF4* allele, we imputed coverage-adjusted IRR count from other haplotypes in the reference panel. In UKB, we selected reference haplotypes based on IBD matches that we had previously computed [11], choosing the top 10 longest IBD matches to heterozygous long-allele carriers in the reference panel (and using all such IBD matches if fewer than 10 were available). We then computed the weighted mean of these reference individuals' coverage-adjusted IRR counts (capped at 25), using weights proportional to $\exp(-c/(\text{IBD length}))$ in order to prioritize more recent IBD-sharing. We tuned the $c$ parameter to maximize the correlation between an individual's own coverage-adjusted IRR and the imputed quantity, selecting $c = 8$ cM in UKB. In AoU, which contained much less IBD-sharing than UKB, we used IBS length instead of IBD and selected $c = 2$ cM.

As noted above, this imputation approach provided imperfect estimates of inherited allele lengths for multiple reasons, including imputation error (arising from germline mutation of the *TCF4* repeat), sampling noise in IRR counts of reference individuals, and upward bias caused by somatic mutation of inherited alleles in reference individuals. However, even rough estimates of relative lengths of inherited *TCF4* alleles were still useful for exploring first-order effects of inherited allele length on somatic mutability (e.g., by stratifying estimated lengths of long *TCF4* alleles (in heterozygotes in the reference panel) by inherited allele length quantile and age; Fig. 3A) and allowed us to partially control for these effects in downstream association analyses.

#### 3.3 GWAS on somatic expansion of long *TCF4* repeat alleles

##### 3.3.1 Optimizing a *TCF4* somatic-expansion phenotype for GWAS

**Quantifying evidence of highly expanded *TCF4* alleles from sequencing data.** In light of our observation that counts of IRR pairs in WGS data (indicative of highly expanded alleles; $\sim$100+ repeat units) were particularly age-associated and thus informative of somatic expansion (Fig. 3A,B and Supplementary Fig. 8A,B), we used this metric as the "raw data" forming the basis for a *TCF4* somatic-expansion phenotype. To increase statistical power, we also computed a similar metric using whole-exome sequencing (WES) data, which provided an independent quantification of *TCF4* expansion for most UKB participants (~470,000; ref. [23]). In detail:

- **WGS-derived *TCF4* expansion metric.** Beyond counting IRR pairs, we also assigned a count of 0.5 to individuals with no IRR pairs but with at least one read pair that barely missed the cutoff for being an IRR pair (i.e., one read in the pair qualified as an IRR based on containing ≥45 CAGs, and its mate contained ≥40 CAGs). We considered adjusting this metric for WGS coverage and mean fragment length but observed a negligible impact on downstream GWAS power (which was unsurprising given that sampling noise was the primary source of noise in counts of IRR pairs; Supplementary Fig. 8B), so we did not adjust for these WGS parameters.
- **WES-derived *TCF4* expansion metric.** We computed a similar metric from WES data, though some additional care was required because (i) WES reads were much shorter than WGS reads (76bp versus 151bp, resulting in lower mapping qualities and many IRR pairs being flagged as potential duplicates) and (ii) technical variation in WES sequencing depth profiles. To handle these issues, we first identified WES read pairs for which:
  - both reads mapped to the *TCF4* CAG repeat in GRCh38 (either mapping completely within the CAG region or partially overlapping it),
  - at most 10 base pairs of each read were unmapped,
  - one read aligned on the forward strand and the other on the reverse strand, and
  - SAM flags `0xB00` were excluded (but duplicate reads were allowed).

  Among these read pairs, we counted the numbers of:

  - IRR pairs (defined based on both reads mapping completely within the CAG region)
  - "IRR+flank" read pairs (for which (a) the left read was an IRR and the right read spanned the right endpoint of the CAG repeat, or (b) the left read spanned the left endpoint of the CAG repeat and the right read was an IRR).

  We adjusted each of these counts for WES coverage and for duplicate read rate. We then further normalized the adjusted IRR pair count by using the adjusted IRR+flank count to control for WES batch effects (since the adjusted IRR+flank count is expected to be the same for all long-allele carriers, up to technical variation in WES sequencing depth profiles). To do so, we computed an individual-specific correction factor equal to the median adjusted IRR+flank count among long-allele carriers (as assessed using WGS data) represented among 300 individuals with similar WES depth profiles [24] (as well as the individual being normalized).

We then computed the WES-derived *TCF4* expansion metric by dividing an individual's adjusted IRR pair count by this correction factor.

**Converting *TCF4* expansion metrics and imputed allele length into a somatic-expansion phenotype by predicting age.** The *TCF4* expansion metrics we computed from WGS and WES data, together with the rough estimates of relative lengths of inherited alleles that we obtained from imputation, provided the main pieces of information we needed to assess an individual's level of somatic expansion. However, the optimal way in which to combine these measurements into a GWAS phenotype was not immediately clear. Ideally, we wished to compute an individual's expected genetic liability for somatic expansion based on these observed quantities. However, the precise relationship between IRR pair counts from sequencing data, imputed allele length, and genetic propensity for expansion was presumably a complicated, nonlinear function, such that running GWAS on an IRR pair count metric and including imputed allele length as a covariate would achieve suboptimal power.

We realized that a more-optimal way to utilize these data was to use mean age as a proxy for the extent to which measurements of WGS- or WES-derived *TCF4* expansion and imputed allele length were informative of somatic expansion. The intuition behind this approach is that if people with a given IRR pair count-derived expansion metric and a given imputed allele length tend to skew older (respectively, younger), this tells us that these measurements generally correspond to higher (respectively, lower) levels of somatic expansion.

To implement this approach, we fit a 9-parameter model to predict age from an individual's IRR pair count-derived *TCF4* expansion metric (either WGS-based or WES-based) and imputed allele length (taking the longer of the two imputed lengths in individuals homozygous for a long *TCF4* allele). We parameterized this model to be able to capture the main nonlinearities of this relationship (Supplementary Fig. 8C):

- 2 parameters $z_{\min}, z_{\max}$ determined cropping thresholds for the *TCF4* expansion metric $z$ (e.g., number of IRR pairs) beyond which smaller or larger values of the metric did not provide additional information about somatic expansion
- 3 parameters defined a quadratic function approximating how age varied as a function of imputed allele length among individuals with the smallest value(s) of the *TCF4* expansion metric (specifically, those cropped to $z_{\min}$)
- 3 parameters defined a quadratic function approximating how age varied as a function of imputed allele length and age among individuals with the largest values of the *TCF4* expansion metric (specifically, those cropped to $z_{\max}$)
- 1 parameter defined an interpolation function determining how to interpolate between the above two curves for individuals with intermediate values of the *TCF4* expansion metric: this function had a value of 1 at $z_{\min}$ and 0 at $z_{\max}$, and the single parameter allowed the

interpolation weight to vary nonlinearly in between (via a quadratic function of $z$).

We optimized these 9 parameters using `optim()` in `R` to minimize the mean squared error (MSE) between predicted and actual age. We could then use the fitted model to predict an individual's age (as a proxy for somatic expansion) given their IRR pair count-derived *TCF4* expansion metric (either WGS-based or WES-based) and imputed allele length (longer of two alleles for hom-long individuals).

We fit the model independently for UKB WGS (Supplementary Fig. 8C), UKB WES, and AoU WGS, obtaining qualitatively similar behavior of these three model fits. To combine WGS-based predicted age and WES-based predicted age into a single somatic-expansion phenotype in UKB, we computed a weighted average of the WGS-based and WES-based predicted ages, selecting the weight that produced the largest correlation with actual age ($\approx 71\%$ WES + 29% WGS). For UKB participants with WGS data but no WES data, we used the WGS-based predicted age phenotype alone.

##### 3.3.2 GWAS sample inclusion criteria

To enable robust, well-powered genome-wide association analysis of somatic expansion of *TCF4* repeats in UKB and AoU, we applied the following sample selection filters.

**Long-allele carriers.** We restricted to individuals who carried long *TCF4* alleles ($\geq$45 repeat units) as described in Section 3.2.2, given that only long alleles were sufficiently unstable to contribute meaningfully to GWAS power. Long-allele carriers represented $\sim$8% of individuals of European genetic ancestry and were a few times less common in non-European ancestries.

**Age and relatedness.** In UKB, we did not apply any filters on age (which ranged from 40–70 years old at DNA acquisition) or relatedness (because we performed GWAS in UKB using a linear mixed model, which controlled for relatedness). In AoU, which had a wider age range (18–90+ years), we restricted to individuals of age $\geq$40, and we further restricted to a subset of unrelated individuals by removing the younger individual in each pair of relatives with kinship coefficient $>$0.1.

**Genetic ancestry.** In UKB, we inferred the genetic ancestry of each participant using their coordinates along the top 20 genetic principal components (PCs). We defined EUR, SAS, AFR, and EAS cluster centers by taking mean PC coordinates among individuals with self-reported ethnic group "White," "Asian or Asian British," "Black or Black British," and "Chinese," respectively, given that self-reported ethnic groups in UKB were previously observed to broadly match the genetic ancestries of 1000 Genomes continental population groups [10]. We then assigned individuals to genetic ancestry clusters based on their Euclidean distances (in PC-space) to these cluster centers. Specifically, for each cluster, we examined Euclidean distances from that cluster center among individuals of the corresponding ethnic group, and we defined a cluster radius that included 99% (for EUR) or 90% (for SAS, AFR, and EAS) of these individuals. We then used these PC-based cluster definitions to assign individuals to genetic ancestry clusters (irrespective of their reported ethnic groups), leaving individuals who did not fall within any cluster unassigned.

In AoU, we used genetic ancestry assignments that had previously been computed [25].

To safeguard against confounding from population stratification, we restricted GWAS to individuals of EUR genetic ancestry in UKB and AoU. We also performed a version of the GWAS that lifted this restriction (instead including genetic ancestry as a covariate) and observed no appreciable difference in the results, which was unsurprising given that the large majority of long-allele carriers in the combined data set (>90%) were of EUR genetic ancestry.

The filters above resulted in GWAS sample sets of size n=40,231 in UKB and n=8,217 in AoU. In addition to these filters, we applied a final sample exclusion criterion that varied based on the genetic variant being tested for association with somatic expansion.

**Chromosome-specific exclusions of individuals with long CAG repeats at loci other than *TCF4*.** Counts of IRR pairs (on which our *TCF4* somatic-expansion phenotype was based) primarily reflected repeat length variation at *TCF4*, but in individuals with long CAG repeat alleles at both *TCF4* and another locus, IRR pairs could sometimes originate from a locus other than *TCF4*. If such individuals were included in a GWAS of somatic expansion at *TCF4*, false positives would arise from variants tagging highly expanded alleles at other loci. To remove such effects while minimally impacting GWAS sample size, we applied a final, chromosome-specific sample exclusion filter: when testing variants on a given chromosome, we excluded individuals with anchored IRRs to loci on that chromosome (other than *TCF4*). We made one exception to this rule: we did not exclude individuals with anchored IRRs at *CA10*, as long *CA10* alleles are common but highly expanded *CA10* alleles are rare (Fig. 1B).

##### 3.3.3 GWAS meta-analysis of somatic repeat expansion in *TCF4*

We ran genome-wide association analysis independently in UKB (on our combined WGS+WES-derived *TCF4* somatic-expansion phenotype) and in AoU (on our WGS-derived somatic-expansion phenotype) and then meta-analyzed the results.

In UKB, we adjusted for the following covariates: sex, age, age$^2$, 20 PCs, imputed allele length, age $\times$ imputed allele length, age$^2$ $\times$ imputed allele length, and coverage-adjusted *CA10* IRR count (i.e., $(\#_{\text{anchored IRRs at } CA10})/(\text{cov}_{\text{GC67}}/\overline{\text{cov}_{\text{GC67}}})$, which captured length variation in *CA10* alleles). We tested TOPMed-imputed variants [26] with MAF>0.1% using BOLT-LMM with the flag `--lmmForceNonInf` to require use of a non-infinitesimal linear mixed model [27,28].

In AoU, we adjusted for the following covariates: sex, age (at DNA acquisition; i.e., "biosample age"), age$^2$, 16 PCs, imputed allele length, age $\times$ imputed allele length, and age$^2$ $\times$ imputed allele length. We tested variants in the ACAF threshold srWGS joint callset (short-read WGS SNP and indel variants with population-specific allele frequency $>$1% or population-specific allele count $>$100 in any ancestry) using linear regression as implemented in BOLT-LMM.

We then meta-analyzed GWAS results from UKB and AoU using METAL [29]. To account for the additional power provided in the UKB GWAS by BOLT-LMM's non-infinitesimal mixed model, we computed standard errors of effect size estimates as $|\beta|/\sqrt{\chi^2_{\text{non-inf}}}$ (as $\beta$ and standard errors in the BOLT-LMM output are from the infinitesimal mixed model). To account for the different scale of the GWAS phenotypes (i.e., predicted age) in UKB versus AoU, we additionally scaled effect sizes and standard errors by the standard deviation of the GWAS phenotype within each cohort. We then performed meta-analysis, weighting according to these updated standard errors, and restricting to variants with minor allele frequency $>$1% in both cohorts.

Finally, we extended the meta-analysis to low-frequency coding variants at loci identified from the common-variant GWAS meta-analysis. Specifically, we identified 67 coding variants with a high or moderate impact VEP score [30] according to the AoU variant annotation table that were within 250kb of a lead variant (Table 1) and that had MAF between 0.1% and 2%. Four such variants associated with *TCF4* somatic expansion at a Bonferroni-significance level of $p$<0.05/67: two missense variants in *FAN1* that have previously been associated with Huntington's disease age-at-onset [31] (chr15:30910758 G>A: *FAN1* Arg507His, $p = 6.6 \times 10^{-11}$; and chr15:30905792 C>T: *FAN1* Arg377Trp, $p = 1.0 \times 10^{-5}$), a missense variant in *MSH3* (chr5:80813660 T>G: *MSH3* Leu911Trp, $p = 5.7 \times 10^{-6}$), and a missense variant in *OCM* that probably tags *PMS2*-related variation (chr7:5882598 G>T, $p = 1.7 \times 10^{-4}$).

##### 3.3.4 Local genetic correlation of somatic repeat expansion in *TCF4* and HD age-at-landmark phenotypes

We computed local genetic correlations between our *TCF4* somatic-expansion phenotype and HD age-at-landmark phenotypes at *MSH3* (hg19: 5:79413275-80413275), *PMS2* (hg19: 7:5522626-6522626), and *FAN1* (hg19: 15:30702961-31702961) using LAVA [32], supplying the provided 1000 Genomes EUR reference data for LD estimation. We lifted summary statistics from our *TCF4* somatic-expansion GWAS to hg19 and estimated local genetic correlations with *hastening of* age-at-SDMT30.

### 4 Phenotypic associations of long CAG repeats

We tested long repeats for association with 57 heritable quantitative traits (including anthropometric traits, blood pressure, measures of lung function, bone mineral density, blood cell indices, and serum biomarkers) [33] and 1,129 health outcomes (organized into ICD-10 categories) in the "first occurrence" data fields that UKB generated by aggregating information from self-report, inpatient hospital data, primary care or death record data. We restricted to n=421,356 unrelated individuals of EUR genetic ancestry (dropping one individual per second-degree or closer relationship pair, i.e., kinship >0.0884 as computed by UKB [10]). We analyzed 14 of the 15 CAG repeat loci that we studied, dropping the repeat locus at chr4:30.7Mb as all but one carrier of a long repeat had non-EUR genetic ancestry.

For each of the 14 repeats, we tested expansion status for association with each of the 57 heritable quantitative traits using linear regression, adjusting for sex, age, and age$^2$ as covariates. Associations that reached a significance threshold of $p < 5\times10^{-5}$ (slightly conservative for $14\times57$ tests) are reported in Supplementary Table 6. We likewise tested expansion status of each repeat for association with each of the 1,129 ICD10 "first occurrence" binary disease phenotypes (accessed on 2022-10-26) using the BinomiRare test [34] to obtain $p$-values robust to case-control imbalance while adjusting for age and sex. As previously described [11,33,35], for computational efficiency, we reimplemented the BinomiRare test and applied a binomial approximation when the number of observed cases among carriers exceeded 100. Associations that reached a significance threshold of $p < 1 \times 10^{-6}$ (conservative for $14 \times 1129$ tests) are reported in Supplementary Table 7.

#### 4.1 Detailed analysis of the *GLS* locus

We investigated the *GLS* 5' UTR CAG repeat locus in more detail in light of the associations that we identified with quantitative and binary traits (Supplementary Tables 6 and 7) as well the role of highly expanded *GLS* repeats in autosomal recessive glutaminase deficiency with impaired intellectual development and progressive ataxia [36].

In these follow-up analyses, we subdivided long *GLS* repeat alleles ($\geq$45 repeat units) into those that were highly expanded ($\sim$100+ repeats, based on the presence of $\geq$1 IRR pair) and those that were not, and we also examined SNPs and indels predicted to cause loss of function (pLoF). We identified carriers of *GLS* pLoF variants using the UKB WES genotype calls [23], restricting to variants annotated as pLoF by gnomAD v4.0.0 [37] with no QC flags (`lc_lof` or `lof_flag`). We additionally filtered two indel variants (`chr2:190931597:C:CA_C` and `chr2:190931597:CA:C_CA`) that had much higher frequences than expected in the UKB WES genotype calls.

We tested each of these three classes of *GLS* variants (highly expanded repeats, long but not highly expanded repeats, and pLoF variants) for association with the 57 heritable quantitative traits, and we also tested for association with glutamine levels measured in plasma (Fig. 5C). Whereas all three classes of variants associated with increased glutamine levels and decreased height with similar effect sizes, only highly expanded repeats associated with liver and renal biomarkers. Associations with highly expanded repeat status that reached Bonferroni significance ($p < 0.05/57$)

are reported in Supplementary Table 8.

For each individual with a long *GLS* allele, we estimated *GLS* allele lengths as described previously (equation (3) from section 3.2.1), assuming any IRR pairs originated from the *GLS* locus. We restricted to individuals with EUR genetic ancestry, removing related individuals among the set of individuals with long *GLS* alleles. We then residualized standardized GGT, cystatin C, and glutamine for sex, age, and age squared. For quintiles of long *GLS* allele lengths, we computed the mean residualized phenotype and mean allele length. We also computed the mean resiudalized phenotype among individuals with no long *GLS* allele. Confidence intervals were computed using a $t$-test.

Given the large effects of highly expanded *GLS* repeat alleles on both liver and renal biomarkers, we tested highly expanded repeat status for association with ICD-10 categories and subcategories of liver diseases (K70–K77, Diseases of liver) and common kidney diseases (N17–N19, Acute kidney failure and chronic kidney disease) using the BinomiRare test. We obtained ICD-10 codes containing up to four characters from hospital episode statistics, cancer registry, and death registry data for UKB participants (accessed on 2022-10-26), from which we compiled 11 category-level ICD-10 disease phenotypes and 62 subcategory-level phenotypes, for a total of 73 liver and renal disease phenotypes.

Among the 11 association tests with top-level ICD-10 categories, three were significant (FDR-adjusted $p < 0.05$), and among the 62 association tests with ICD-10 subcategories, four were significant (FDR-adjusted $p < 0.05$); results are reported in Supplementary Table 9. The strongest association we observed was with the ICD-10 subcategory for stage 5 chronic kidney disease (N18.5). Upon closer examination of ICD-10 subcategory definitions within N18, we found that a separate code is used for stage 5 CKD requiring chronic dialysis (N18.6, or previously N18.0, for "end-stage renal disease"), such that combining these subcategories would provide a better phenotype for stage 5 CKD. We therefore focused on this phenotype (CKD stage 5, inclusive of end-stage renal disease) in our primary analyses.

#### 4.2 Replication of *GLS* associations with liver and kidney disease in *All of Us*

We identified carriers of highly expanded *GLS* repeats in the AoU v7 WGS data set based on the presence of an anchored IRR at *GLS* and at least one IRR pair. We then attempted to replicate associations we observed in UKB between highly expanded repeat status and the following liver and kidney disease phenotypes:

- "Other diseases of liver" (K76), defined as an ICD-10 code of K76 or an ICD-9 code of 573.0, 572.3, 572.4, or 573.5 (corresponding to the ICD-10 subcategories of K76)

- Chronic kidney disease (N18), defined as the "Chronic kidney disease" concept in AoU (which aggregates both ICD-10 and ICD-9 codes)
- CKD stage 5, inclusive of end-stage renal disease, defined as an ICD-10 code of N18.5 or N18.6 or an ICD-9 code of 585.5 or 585.6.

We did not attempt to replicate the association with the ICD-10 top-level category K75 ("Other inflammatory liver diseases") as power was very limited due to low case prevalence in AoU, and similarly, we did not attempt to replicate associations with ICD-10 subcategories other than stage 5 CKD.

We performed association tests using logistic regression, adjusting for sex, age, $age^2$, and genetic ancestry, and restricting to an unrelated subset of individuals (removing one individual in each pair of relatives with kinship coefficient $>0.1$, retaining carriers of highly expanded *GLS* alleles when possible).

### 5 Supplementary Figures

**A**

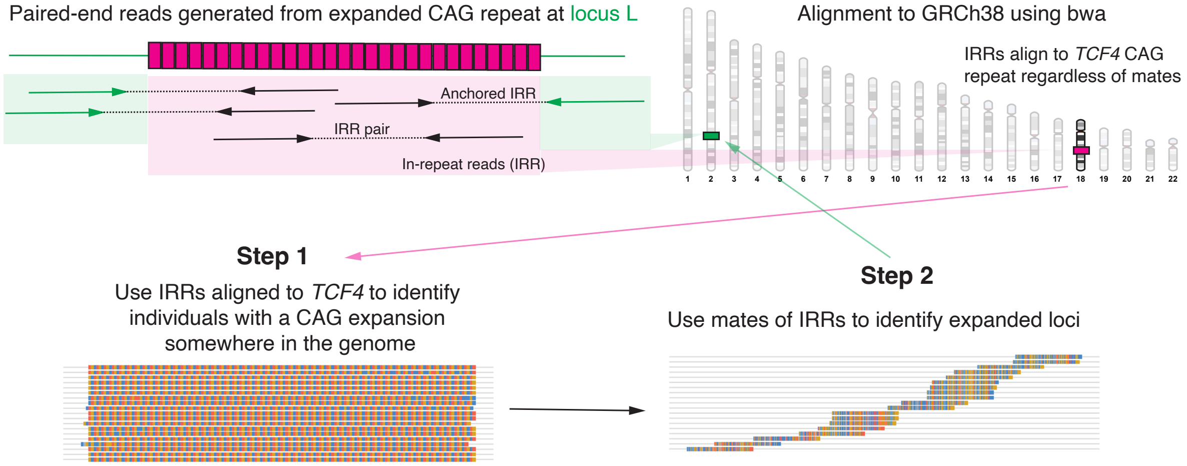

**B**

| | Number of IRRs observed among 500 random WGS samples | | |
|---|---|---|---|
| Occurrences of repeat motif in read SEQ | IRRs mapped to *TCF4* | IRRs mapped elsewhere with mates mapped to *CA10* | Other IRRs |
| 45 | 84 | 38 | 9 |
| 46 | 94 | 20 | 4 |
| 47 | 115 | 7 | 5 |
| 48 | 132 | 11 | 4 |
| 49 | 170 | 10 | 7 |
| 50 | 643 | 5 | 0 |
| 45-50 | 1238 | 91 | 29 |

**C**

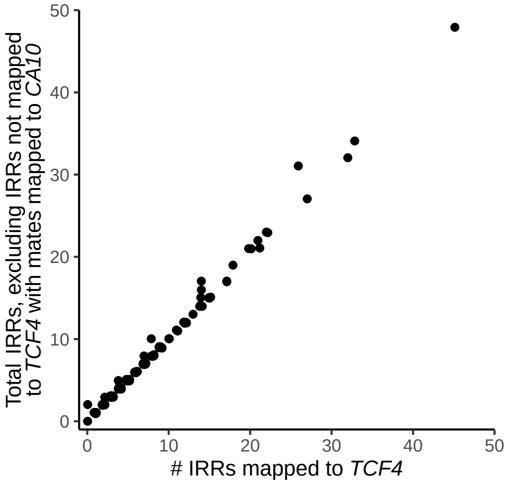

**Supplementary Figure 1. Overview of approach to identify long CAG repeats from short-read WGS read alignments. A** Individuals with long CAG repeat alleles have WGS reads consisting entirely or almost entirely of CAG repeats (in-repeat reads, IRRs). The bwa aligner [1] maps most such reads to the CAG repeat in *TCF4* on chromosome 18 (which is the longest CAG repeat in the GRCh38 reference genome), such that searching for IRRs among reads aligned to *TCF4* reveals which individuals have long CAG repeats somewhere in their genome. By examining the mapping locations of uniquely-mapped mates of IRRs ("anchoring" these IRRs), we can determine where in the genome these repeat sequences originated. **B** Mapping of IRRs found across 500 randomly selected WGS cram files (of which 164 had at least one IRR). Each IRR is classified as either (i) mapping to *TCF4*, (ii) not mapping to *TCF4* but with a mate that mapped to *CA10*, or (iii) not mapping to *TCF4* and with a mate that mapped to locus other than *CA10*. **C** The number of IRRs per individual satisfying (i) or (iii) is plotted versus the number satisfying (i).

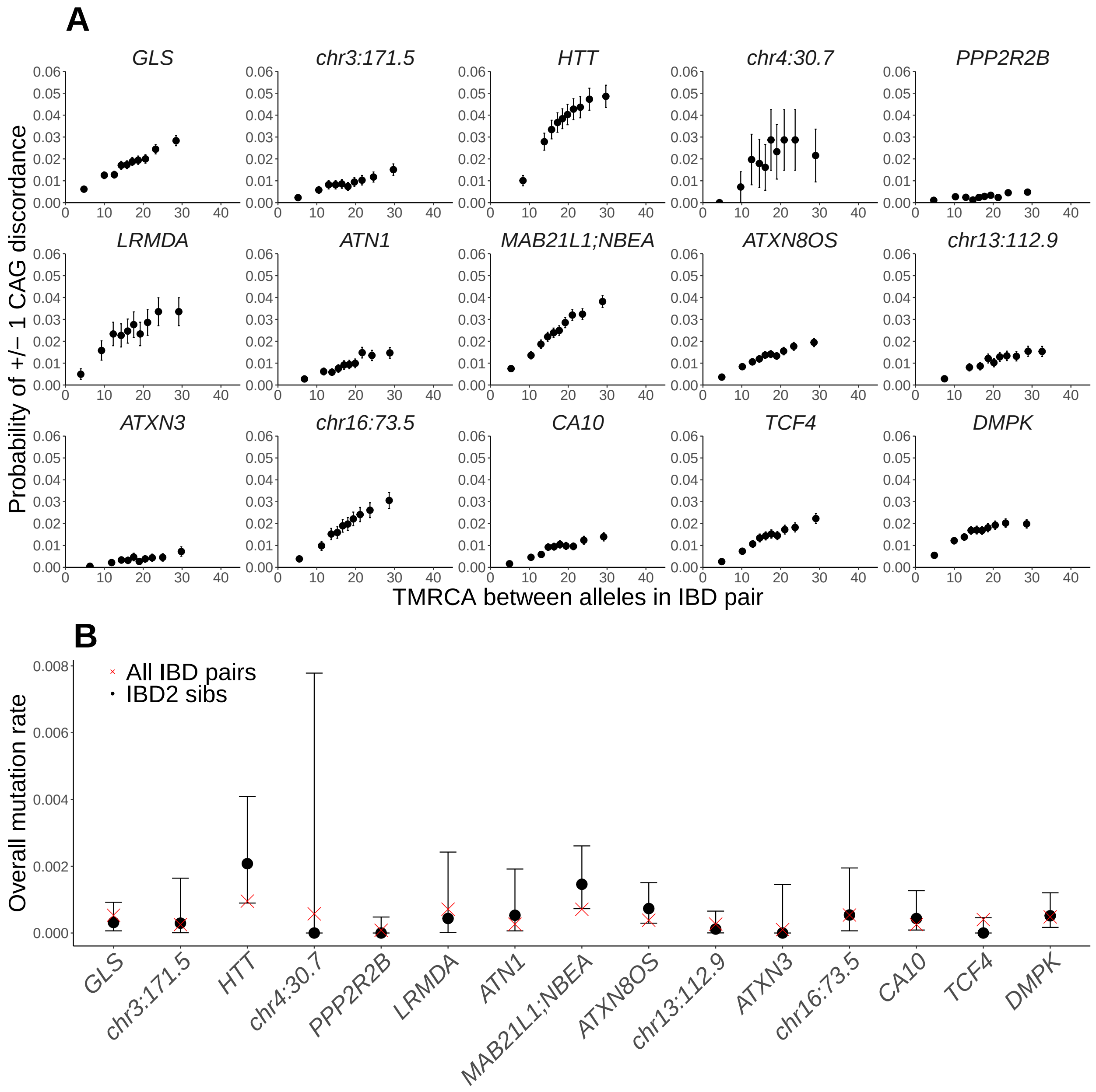

**Supplementary Figure 2. Validation of germline mutation rates of CAG repeats.**
**A** Discordance rate among alleles shared within IBD segments as a function of estimated TMRCA (binned into deciles). Discordance rates scale approximately linearly with TMRCA, as expected if discordances mostly reflect recent germline mutations and if TMRCA is accurately estimated. **B** Rates of $\pm 1$ CAG mutations per transmission of each repeat directly estimated from IBD2 sibling pairs (black dots; error bars, 95% CIs) and estimated from all IBD pairs used in our analyses of allele-specific germline mutation rates (red crosses).

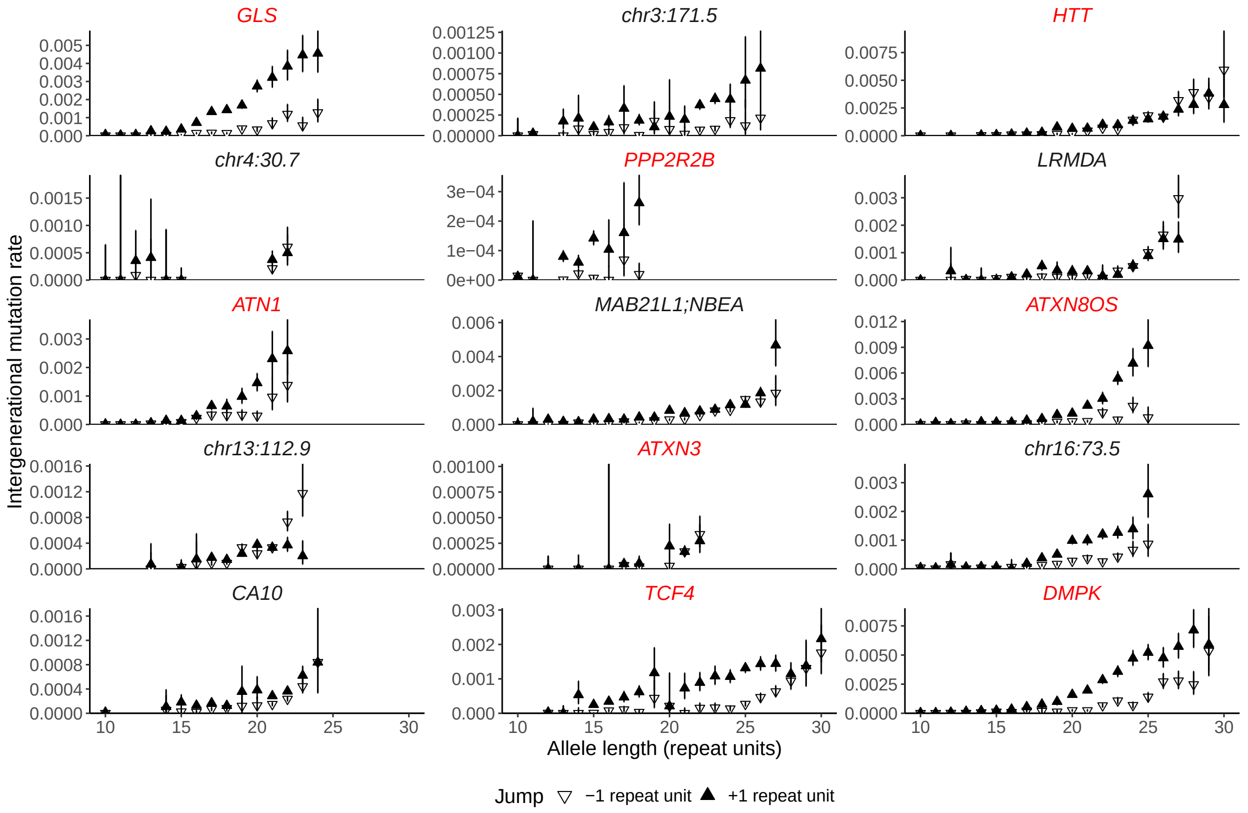

**Supplementary Figure 3. Germline mutation rates of CAG repeats.** Estimated per-generation mutation rates of germline expansion (+1 repeat unit) and contraction (–1 repeat unit) of 15 repeat loci in UKB. Repeat expansions in genes highlighted in red are known to be pathogenic.

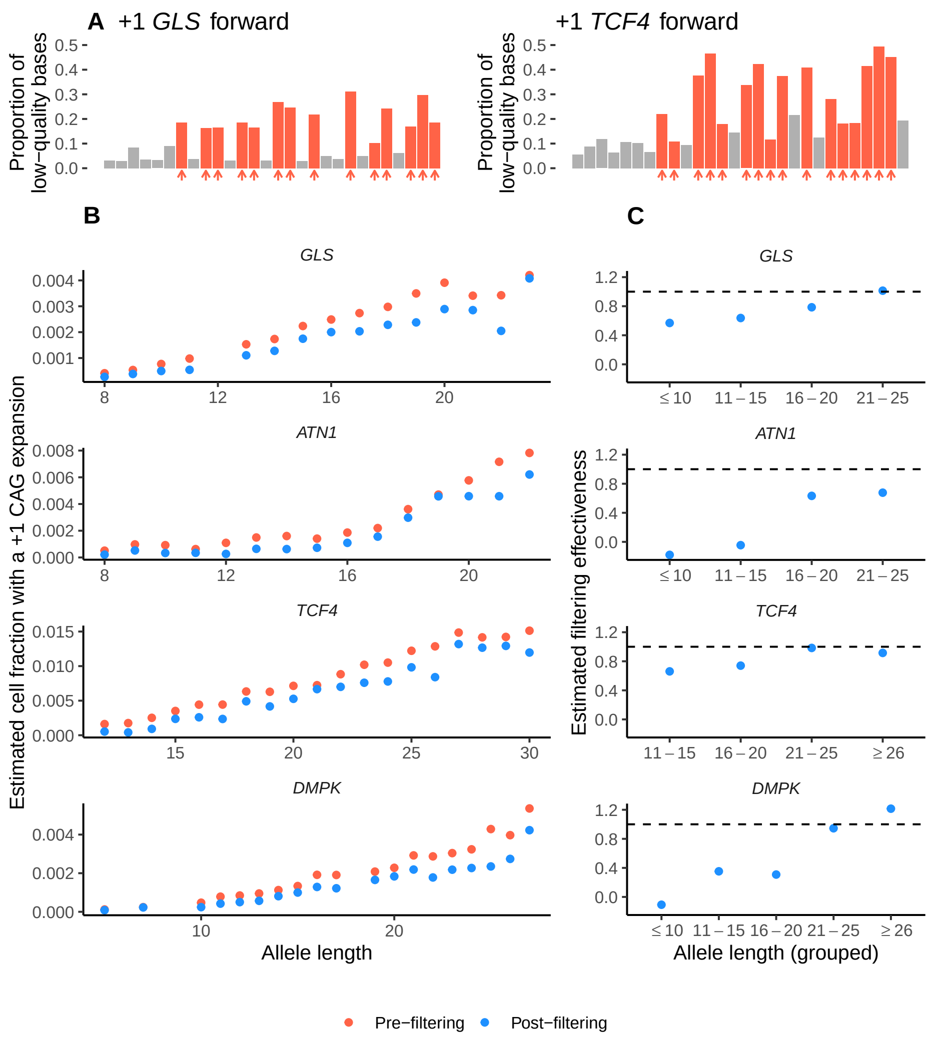

**Supplementary Figure 4. Effect on somatic instability analysis of filtering aberrant reads with evidence of PCR stutter error. A** PCR stutter errors during bridge amplification generate "barcodes" of reduced-quality base calls at positions following the repeat at which the original sequence differs from the expanded/contracted sequence (see Fig. 2D). For *GLS* (left) and *TCF4* (right), proportions of low-quality bases among UKB WGS forward-strand reads supporting a +1 repeat unit expansion (some truly derived from somatic mutations, others generated by PCR error) are shown at the final six base positions of the repeat and positions thereafter. Bases at which PCR stutter error is predicted to reduce base quality are indicated with red arrows.
**B** Estimated fractions of blood cells in which UKB participants heterozygous for a repeat allele of a given length had experienced a +1 repeat unit expansion, computed either using all reads (pre-filtering) or applying our filtering approach. **C** Estimated filtering effectiveness (i.e., proportion of reads passing filtration that were truly derived from somatically-expanded +1 repeat alleles) as a function of allele length. These estimates were obtained by regressing the fraction of cells estimated to have a +1 repeat unit expansion on age in AoU.

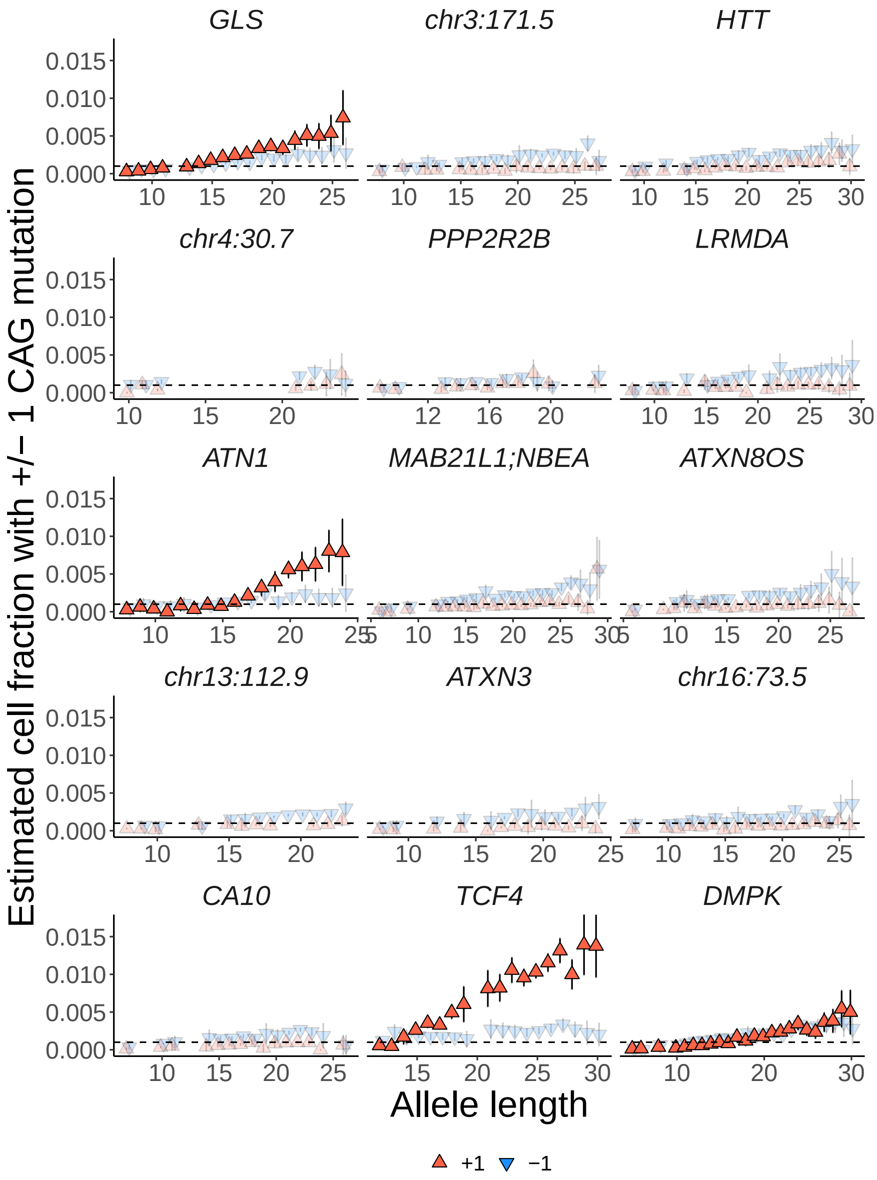

**Supplementary Figure 5. Estimated fraction of blood cells with a ±1 repeat unit mutation as a function of allele length.** For each repeat locus, frequencies of somatic +1 unit expansions and −1 unit contractions were estimated among all UKB participants heterozygous for a repeat allele of a given length. For most repeat loci, estimated contraction frequencies exceeded estimated expansion frequencies and these estimates did not significantly increase with age, indicating that even after applying our filtering approach, the estimates were still largely reflecting residual PCR stutter error rather than somatic mutation. For four repeat loci (*GLS*, *ATN1*, *TCF4*, and *DMPK*), estimated frequencies of +1 unit expansions associated significantly with age; these estimates are indicated with darker markers.

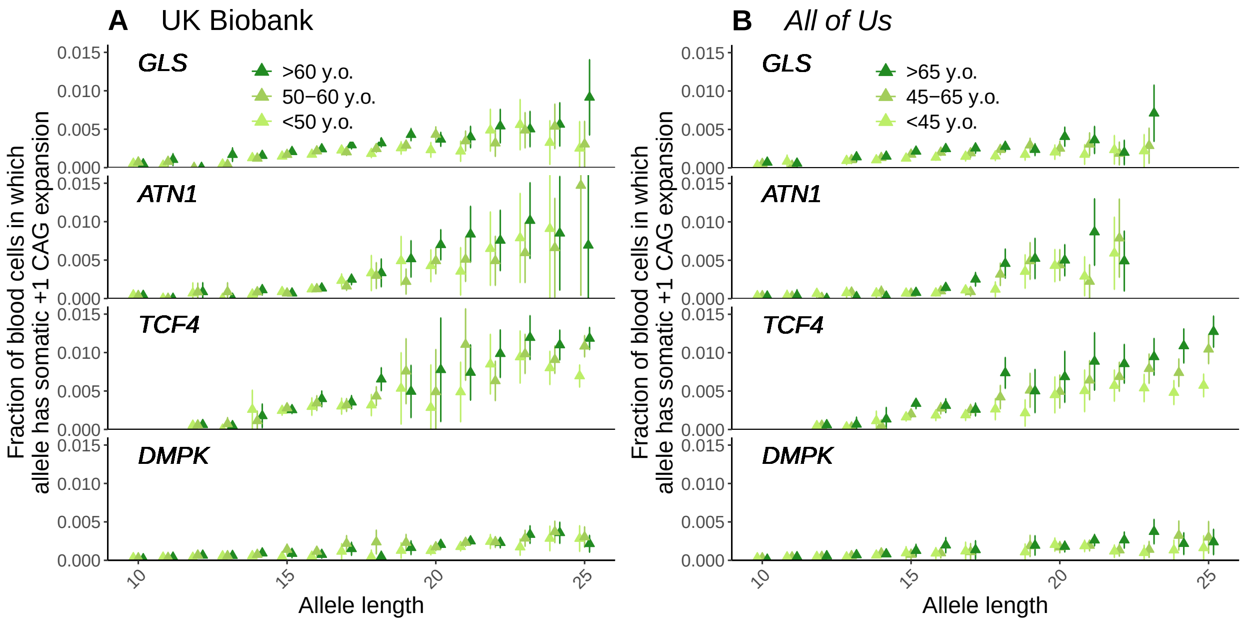

**Supplementary Figure 6. Estimated fractions of blood cells with +1 repeat unit expansions increase with age for four CAG repeat loci.** Estimated frequencies of somatic +1 unit expansions among individuals heterozygous for a repeat allele of a given length are shown for three age strata of UK Biobank (**A**) and *All of Us* (**B**).

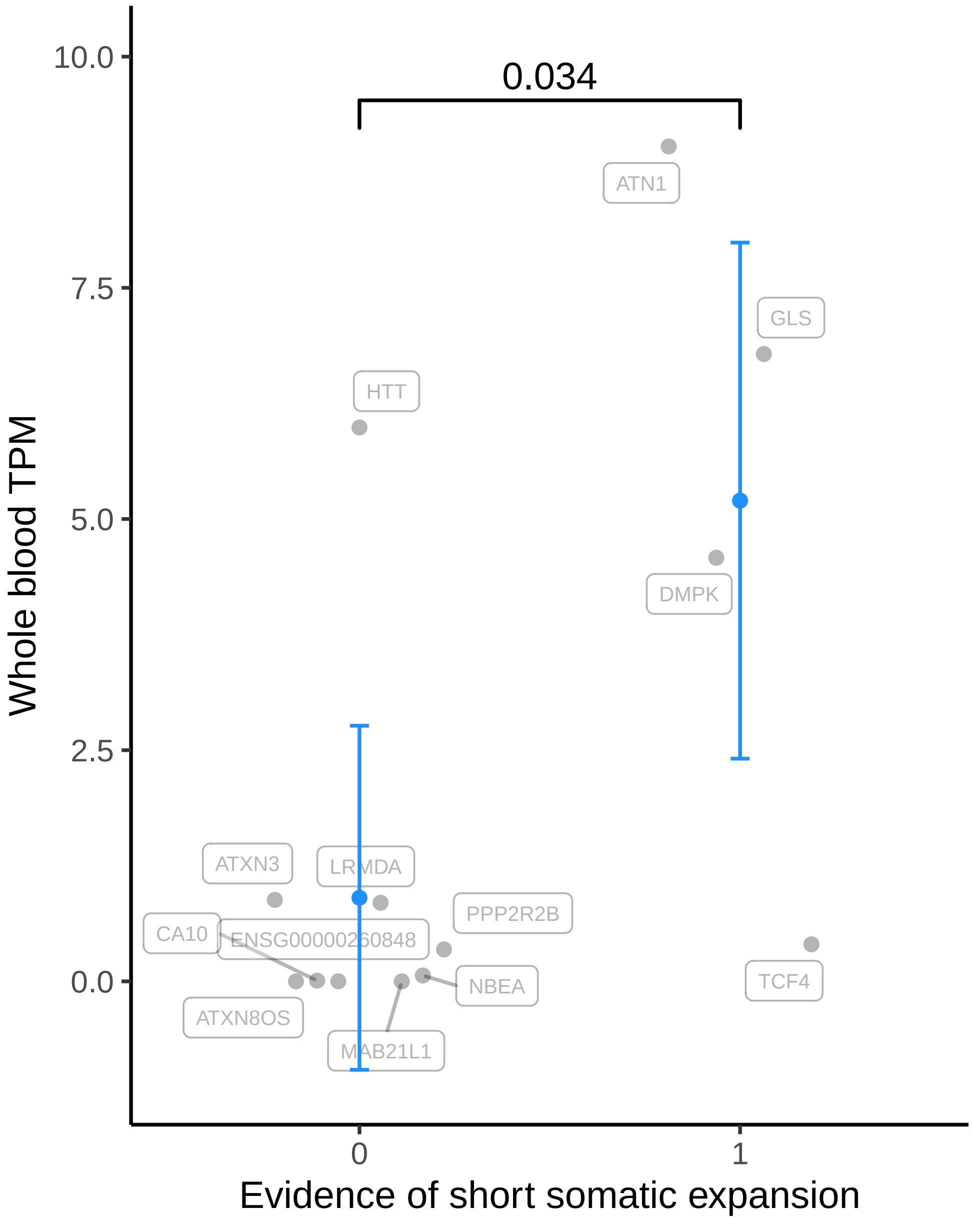

**Supplementary Figure 7. Transcription levels in blood for genes for which repeat expansions are (or are not) expansion-prone in blood.** For each of the 13 genes containing a repeat included in our analyses (Fig. 1B), we obtained its median expression in blood (transcripts per million; TPM) from GTEx v8 [38]. We then classified genes based on whether or not we had identified evidence of somatic repeat instability in blood (Supplementary Figure 5). Each group's mean transcription level (and 95% confidence interval) is shown in blue; p-value for the difference between the groups was computed using the Wilcoxon rank-sum test and shown above.

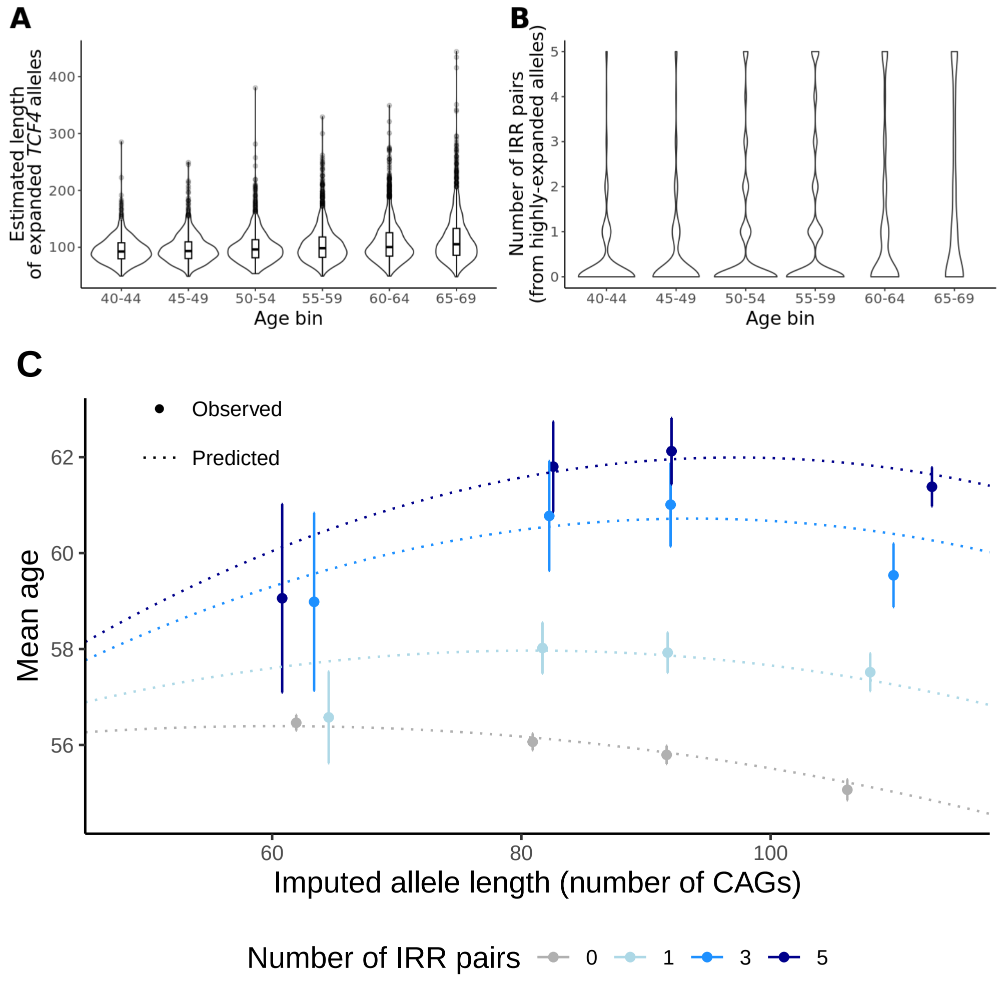

**Supplementary Figure 8. Distributions of *TCF4* allele length metrics and relationship between age, imputed *TCF4* allele length, and count of IRR pairs.** **A**, **B** Distributions of individual-level measurements of estimated lengths of long (≥45-repeat) *TCF4* alleles (panel **A**) and counts of IRR pairs observed in UKB WGS data (panel **B**, capped at 5) in 5-year age tranches of individuals in the highest quintile of imputed allele length (a metric that captures relative lengths of inherited alleles). While there is a strong statistical signal that these length metrics increase with age in UKB (Fig. 3A,B), the individual-level data are noisy and only weakly informative of somatic expansion. **C** Mean age (error bars, 95% CIs) among individuals stratified by IRR pair count and by imputed allele length ($x$-values of dots are means per quartile). For each stratum of IRR pair count shown in the figure (count = 0, 1, 3, 5), mean age predicted by our model is plotted as a function of imputed allele length (dotted lines). On the $x$-axis, imputed allele lengths are computed from imputed coverage-adjusted counts of anchored IRRs at *TCF4* using the conversion $45 +$ (imputed # IRRs) $\times$ $424/$(mean coverage in UKB $\times$ $3$), as derived in equation (3).

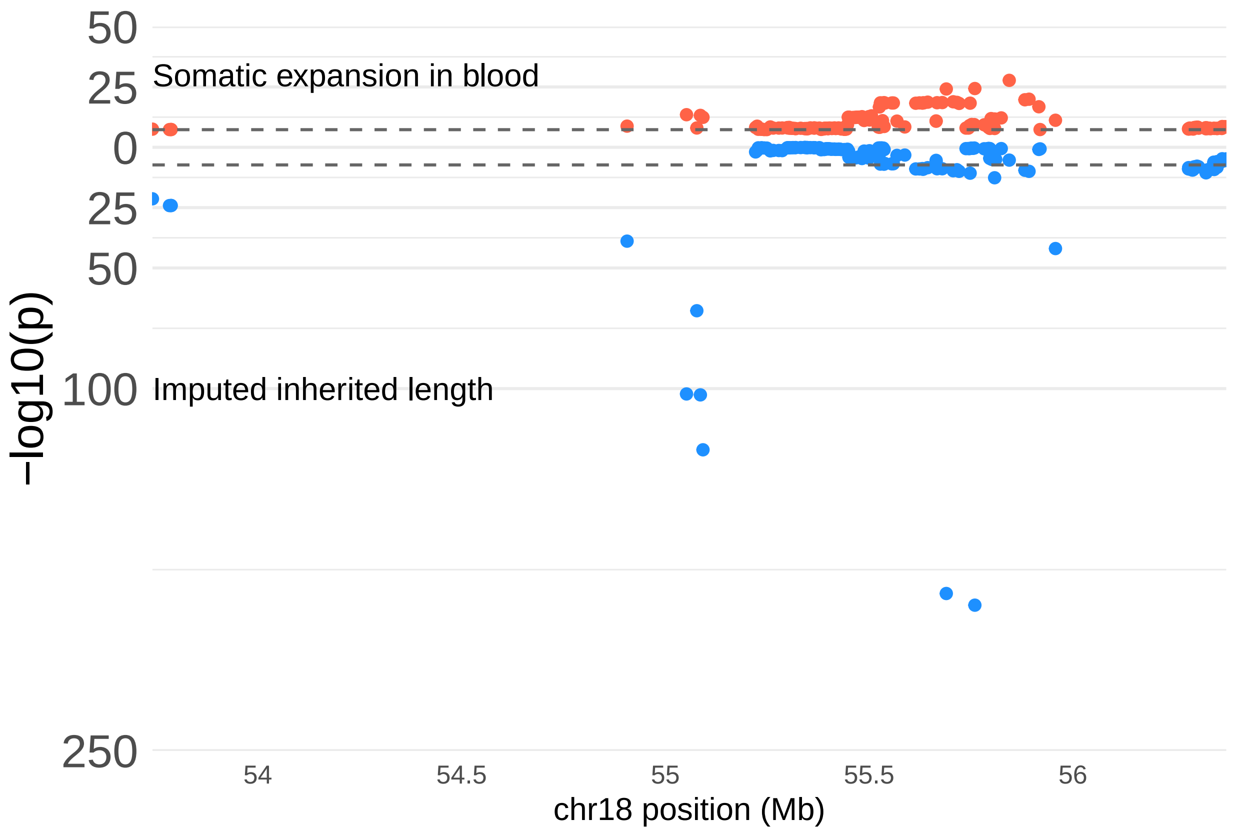

**Supplementary Figure 9. Variants at *TCF4* that associate with somatic expansion of *TCF4* repeats probably tag long inherited alleles.** The top of the plot (red dots) shows associations with *TCF4* somatic expansion in blood; the bottom of the plot (blue dots) shows associations with imputed allele length. Only variants with MAF$>1\%$ that associated with *TCF4* somatic expansion ($p < 5 \times 10^{-8}$) are shown. Association tests were performed using linear regression analyses of long-allele carriers in UKB. In the top analysis, we used the same set of covariates as in our *TCF4* somatic-expansion GWAS. These covariates included imputed allele length (as well as interactions of imputed allele length with age). However, we suspected that these covariates incompletely controlled for the contribution of inherited *TCF4* allele length to our GWAS phenotype, allowing some variants at *TCF4* that tag long inherited alleles to still reach significance in the GWAS (despite having no effect on somatic expansion). The bottom plot supports this interpretation: the variants at *TCF4* that associated with the *TCF4* somatic-expansion phenotype show a similar association pattern with imputed allele length (which captures only inherited allele length variation and does not capture somatic expansion).

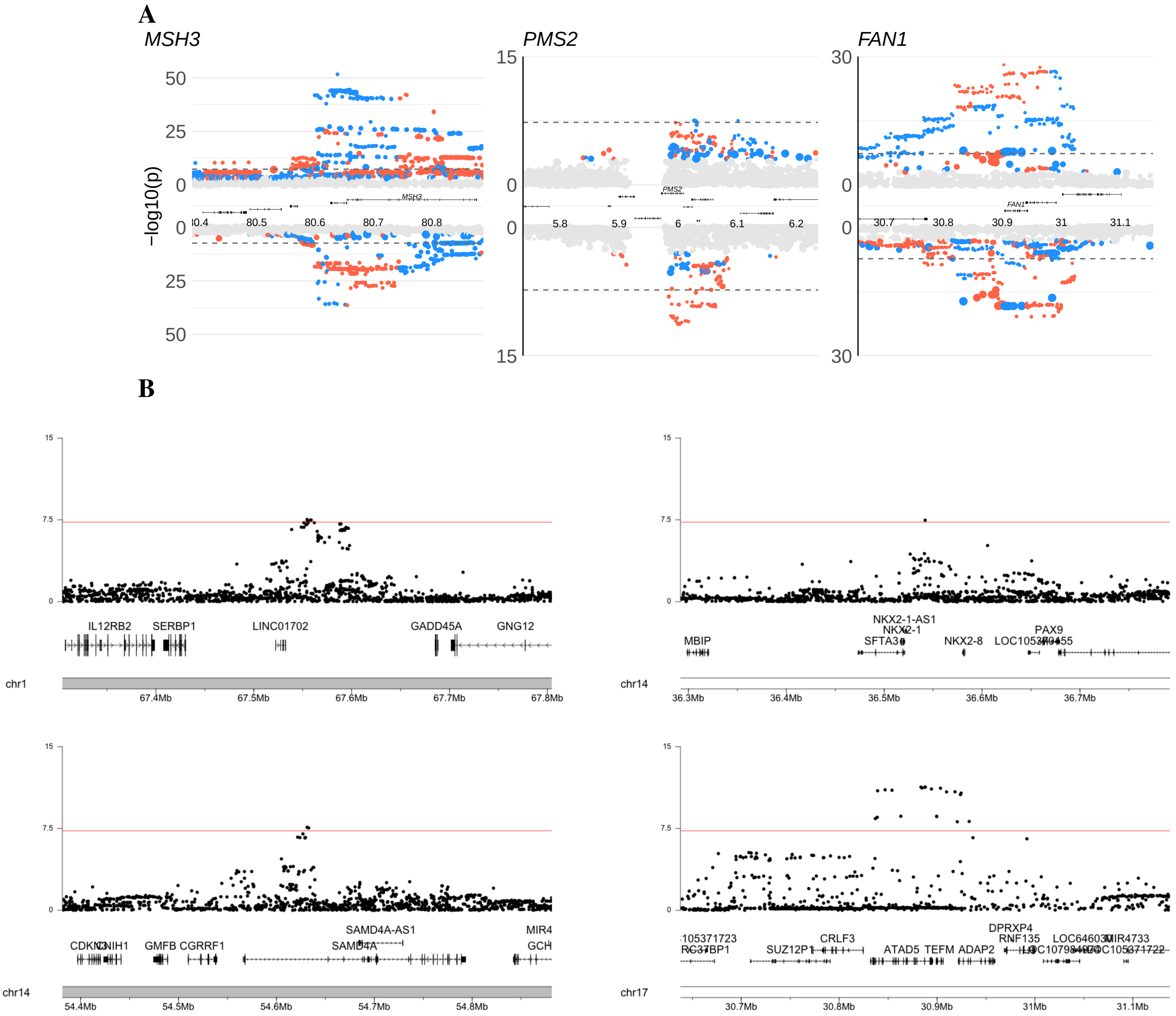

**Supplementary Figure 10. Locus plots of genetic associations with somatic expansion of *TCF4* repeats in blood. A** For loci previously associated with Huntington's disease age-at-onset and age-at-landmark phenotypes (*MSH3*, *PMS2*, and *FAN1*), "Miami" plots show associations with somatic expansion of *TCF4* in blood from our meta-analysis of UKB+AoU (top) and previously-reported associations with age-at-SDMT30 [39] (bottom). Marker colors indicate effect directions for the alternative allele, with blue dots corresponding to expansion-accelerating effects (i.e., increased *TCF4* somatic expansion (top) and hastened age-at SDMT30 (bottom)) and red dots corresponding to expansion-slowing effects. Larger dots indicate rarer variants (i.e., lower minor allele frequency). **B** Associations with somatic expansion of *TCF4* in blood for each of the other four loci that reached significance in our meta-analysis of UKB+AoU. Plots were generated using the karyoploteR [40] package in R [14].

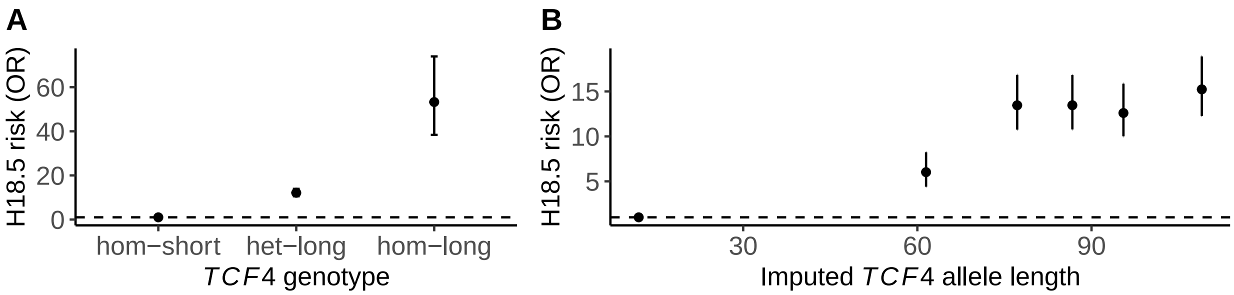

**Supplementary Figure 11. Risk of hereditary corneal dystrophies (H18.5, under which FECD is classified) conferred by long *TCF4* repeats. A** Risk of H18.5 among individuals heterozygous (respectively, homozygous) for a long *TCF4* allele relative to individuals who do not carry a long *TCF4* allele. **B** Risk of H18.5 among individuals heterozygous for a long *TCF4* allele, stratified into quintiles of imputed *TCF4* allele length (a proxy for inherited allele length). Odds ratios were estimated in UKB participants of EUR genetic ancestry using logistic regression, controlling for age and sex. Error bars, 95% CIs.

### 6 Supplementary Tables

| Repeat region (GRCh38) | Genic context | Repeat composition | Flanking sequence Start | End |
|---|---|---|---|---|
| chr2:190880873-190880920 | *GLS* | $GCA_n$ | TAGCGCGCA | GCACCCGCA |
| chr3:171524288-171524320 | Intergenic | $CTG_n$ | AGAACACTG | CTGACTCAG |
| chr4:3074877-3074968 | *HTT* | $CAG_n(CAACAG)_{1,2}$ | TCCTTCCAG | CAGCCGCCA |
| chr4:30716894-30716931 | Intergenic | $AGC_n$ | CGCCAAAGC | AGCAGGCTC |
| chr5:146878728-146878759 | *PPP2R2B* | $GCT_n$ | CTCGCAGCT | GCTGCAGGA |
| chr6:16327634-16327724 | *ATXN1* | $TGC_{n_1}$ $(TGATGC)_{n_2}TGA$ $(TGC)_{n_3}$ | CTGAGGTGC | TGCTCAGCC |
| chr10:75973021-75973073 | *LRMDA* | $CAG_n$ | TAACAACAG | CAGCTACTC |
| chr11:93535787-93535815 | *SRP14P2;SMCO4* | $CAG_nCAA_1CAG_2$ | CCAAAGCAG | CAGCACAGT |
| chr12:6936717-6936773 | *ATN1* | $CAGCAACAGCAA(CAG)_n$ | CACCACCAG | CAGCATCAC |
| chr13:35476296-35476355 | *MAB21L1;NBEA* | $CTG_n$ | GTTTCCCTG | CTGCTTTTC |
| chr13:67656824-67656870 | lncRNA | $AGC_n$ | AATTGGAGC | AGCAGGAAA |
| chr13:70139352-70139429 | *ATXN8OS* | $CTG_n$ | CTACTACTG | CTGCATTTT |
| chr13:112934556-112934581 | Intergenic | $TGC_n$ | AGGGGATGC | TGCTGGTCT |
| chr14:92071009-92071053 | *ATXN3* | $(CTG)_n$ $(TTGCTG)_{0,1}CTTTTG(CTG)_2$ | GTCCCCCTG | CTGTCTGAA |
| chr16:73546663-73546741 | lncRNA | $(TGC)_2$ TGT $(TGC)_nTGTTGC$ | TTTGGGTGC | TGCTTCTTT |
| chr17:51831667-51831732 | *CA10* | $AGC_n$ | AGAAAAAGC | AGCAGAAAA |
| chr18:55586154-55586228 | *TCF4* | $AGC_n$ | AGGAGGAGC | AGCATGAAA |
| chr19:45770205-45770264 | *DMPK* | $CAG_n$ | TCCCCCCAG | CAGCATTCC |

**Supplementary Table 1. Locations and sequence contexts of CAG repeat loci expanded to ≥45 repeat units in ≥5 UKB participants.** Our analysis pipeline used anchored IRRs to approximately localize long repeats to their loci of origin, after which we obtained base-pair-resolution information about repeat locations from ref. [3]. The repeat length polymorphism we measured corresponds to the subscript $n$ in the repeat composition column. The *ATXN1* repeat (chr6:16327634–16327724) contains two distinct length-polymorphic CAG sequences, so we dropped this repeat from downstream analysis along with two other repeats that failed other filters, leaving the 15 loci shown in Fig. 1B. Repeat expansions in genes highlighted in red are known to be pathogenic.

| Genic context | # CAG repeat loci | # expanded | OR (95% CI) | $p$-value |
|---|---|---|---|---|
| intergenic | 328 | 3 | 0.5 (0.09-1.79) | 0.43 |
| transcript | 831 | 15 | 1.99 (0.56-10.8) | 0.43 |
| exon | 331 | 10 | 3.19 (1.12-9.39) | 0.016 |
| CDS | 230 | 4 | 1.16 (0.27-3.73) | 0.77 |
| UTR | 73 | 4 | 4.43 (1.03-14.61) | 0.023 |

**Supplementary Table 2. Enrichment of CAG repeats expanded to $\geq$45 repeat units in $\geq$5 UKB participants in various genic contexts.** Odds ratios, confidence intervals, and $p$-values are from Fisher's exact test for association between genic context of a CAG repeat and whether or not the repeat was expanded in $\geq$5 UKB participants).

| Variant (GRCh38) | Locus | ALT allele frequency (UKB) | $p$-value for association with somatic *TCF4* repeat expansion UK Biobank (n=40,231) | *All of Us* (n=8,217) |
|---|---|---|---|---|
| $p < 5 \times 10^{-8}$ in UKB | | | | |
| 5:80638411:G:T | *MSH3* | 0.37 | $1.4 \times 10^{-49}$ | $3.4 \times 10^{-5}$ |
| 7:5995187:A:G | *PMS2* | 0.62 | $3.0 \times 10^{-9}$ | 1.0 |
| 15:30902611:T:C | *FAN1* | 0.49 | $8.5 \times 10^{-29}$ | 0.019 |
| 17:30883577:G:A | *ATAD5* | 0.27 | $2.0 \times 10^{-9}$ | $4.2 \times 10^{-4}$ |
| $p < 10^{-6}$ in UKB | | | | |
| 1:67554471:T:C | *GADD45A* | 0.97 | $3.7 \times 10^{-7}$ | 0.026 |
| 3:14176823:G:A | | 0.49 | $3.8 \times 10^{-7}$ | 0.87 |
| 14:36541884:A:G | *SFTA3,NKX2-1* | 0.07 | $4.1 \times 10^{-7}$ | 0.028 |
| 14:54631346:A:G | *SAMD4A* | 0.74 | $6.6 \times 10^{-7}$ | 0.011 |
| 20:40979553:T:G | | 0.01 | $9.3 \times 10^{-7}$ | 0.68 |

**Supplementary Table 3. Genome-wide significant ($p < 5 \times 10^{-8}$) and suggestive ($p < 10^{-6}$) associations with somatic *TCF4* repeat expansion in UKB and replication in AoU.** We identified common (MAF>0.01) variants that reached genome-wide or suggestive significance in UKB and were also genotyped in the AoU ACAF WGS data set. Variants at nine genomic loci satisfied these criteria; for the lead variant each locus, data for association in UKB and attempted replication in AoU are provided. GWAS sample sizes in UKB and AoU are much smaller than the full cohort sizes primarily due to GWAS being restricted to long-allele carriers of age at least 40.

| | | | | Age at onset | | Age at SDMT30 | | Somatic *TCF4* expansion | |
|---|---|---|---|---|---|---|---|---|---|
| Variant | Gene | Modifier | MAF | $\beta$ | $P$ | $\beta$ | $P$ | $\beta$ | $P$ |
| 2:190579929:A:C | *PMS1* | 2AM1 | 20.63 | -0.11 | $4.9 \times 10^{-9}$ | -0.086 | $6.6 \times 10^{-5}$ | -0.0008 | 0.91 |
| 2:190639862:C:T | *PMS1* | 2AM2 | 1.28 | 0.3 | $3.7 \times 10^{-6}$ | 0.17 | 0.026 | 0.0081 | 0.81 |
| 3:37068079:T:C | *MLH1* | 3AM1 | 31.41 | 0.11 | $3 \times 10^{-11}$ | 0.088 | $4.5 \times 10^{-6}$ | -0.016 | 0.019 |
| 3:37121844:G:A | *MLH1* | 3AM1 | 31.52 | 0.11 | $1.1 \times 10^{-10}$ | 0.09 | $2.5 \times 10^{-6}$ | -0.017 | 0.014 |
| 4:2533199:G:A | *HTT* | CAA-loss | 0.32 | -1.2 | $4.8 \times 10^{-9}$ | -2.1 | $1.8 \times 10^{-9}$ | | |
| 4:2798279:C:T | *HTT* | CAA-loss | 0.36 | -1.2 | $4.9 \times 10^{-9}$ | -2 | $3.1 \times 10^{-10}$ | | |
| 4:2971698:A:G | *HTT* | CAACAG-dup | 0.60 | 0.52 | $1.7 \times 10^{-7}$ | 0.49 | $4.2 \times 10^{-5}$ | -0.11 | 0.18 |
| 5:79913275:G:A | *MSH3* | 5AM1 | 25.44 | -0.12 | $1 \times 10^{-11}$ | -0.25 | $1.4 \times 10^{-36}$ | -0.076 | $3.6 \times 10^{-25}$ |
| 5:79950781:**<u>A</u>**:G | *MSH3* | 5AM1 | 25.36 | -0.12 | $2.6 \times 10^{-11}$ | -0.25 | $4.7 \times 10^{-37}$ | -0.075 | $1.8 \times 10^{-24}$ |
| 5:80086504:A:G | *MSH3* | 5AM2 | 0.32 | 0.74 | $5.7 \times 10^{-9}$ | 0.59 | $2.9 \times 10^{-5}$ | 0.012 | 0.79 |
| 5:79961856:AT:A | *MSH3* | 5AM3 | 27.78 | 0.078 | $5.6 \times 10^{-6}$ | 0.19 | $2 \times 10^{-22}$ | 0.097 | $4.2 \times 10^{-44}$ |
| 5:79990883:T:G | *MSH3* | 5AM3 | 33.14 | 0.084 | $2.6 \times 10^{-7}$ | 0.17 | $1.4 \times 10^{-19}$ | 0.075 | $3.7 \times 10^{-30}$ |
| 5:145886836:G:A | *TCERG1* | 5BM1 | 2.71 | 0.32 | $3.3 \times 10^{-12}$ | 0.21 | $3.8 \times 10^{-5}$ | -0.014 | 0.47 |
| 7:6022626:C:T | *PMS2* | 7AM1 | 14.67 | 0.12 | $6.4 \times 10^{-9}$ | 0.13 | $1.3 \times 10^{-7}$ | -0.02 | 0.04 |
| 7:6056484:G:C | *PMS2* | 7AM2 | 18.41 | -0.099 | $2.4 \times 10^{-7}$ | -0.088 | $6.7 \times 10^{-5}$ | -0.0046 | 0.55 |
| 7:6041836:T:A | *PMS2* | 7AM3 | 41.42 | 0.075 | $8.3 \times 10^{-7}$ | 0.12 | $4.7 \times 10^{-12}$ | -0.03 | $1.5 \times 10^{-6}$ |
| 7:6026530:C:T | *PMS2* | 7AM4 | 2.14 | -0.17 | 0.00064 | -0.27 | $5.5 \times 10^{-6}$ | 0.049 | 0.021 |
| 7:56241504:C:T | *?* | 7BM1 | 2.94 | 0.26 | $7.3 \times 10^{-9}$ | 0.098 | 0.06 | 0.021 | 0.22 |
| 8:103213640:G:T | *RRM2B* | 8AM1 | 7.91 | -0.16 | $2.1 \times 10^{-9}$ | -0.09 | 0.0042 | -0.013 | 0.33 |
| 11:96079307:C:A | *CCDC82* | 11AM1 | 19.53 | 0.096 | $4.2 \times 10^{-7}$ | 0.084 | $9.2 \times 10^{-5}$ | -0.0004 | 0.94 |
| 15:31202961:G:A | *FAN1* | 15AM1 | 1.24 | -0.69 | $3.8 \times 10^{-28}$ | -0.7 | $1.2 \times 10^{-20}$ | 0.2 | $6.6 \times 10^{-11}$ |
| 15:31247852:G:T | *FAN1* | 15AM1 | 1.15 | -0.7 | $1.3 \times 10^{-26}$ | -0.75 | $4.1 \times 10^{-21}$ | 0.23 | $5.2 \times 10^{-13}$ |
| 15:31241346:G:A | *FAN1* | 15AM2 | 27.81 | 0.19 | $7.2 \times 10^{-28}$ | 0.19 | $1.3 \times 10^{-21}$ | 0.053 | $9.7 \times 10^{-15}$ |
| 15:31197995:C:T | *FAN1* | 15AM3 | 0.82 | -0.59 | $1.8 \times 10^{-13}$ | -0.49 | $6.5 \times 10^{-8}$ | 0.16 | $1 \times 10^{-5}$ |
| 15:31204637:C:T | *FAN1* | 15AM5 | 1.88 | -0.33 | $4.5 \times 10^{-10}$ | -0.18 | 0.004 | 0.096 | $1.6 \times 10^{-5}$ |
| 19:48622545:A:G | *LIG1* | 19AM1 | 17.19 | 0.12 | $1.2 \times 10^{-9}$ | 0.078 | 0.00071 | 0.018 | 0.26 |
| 19:48687051:ACC:A | *LIG1* | 19AM1 | 12.60 | 0.11 | $6.2 \times 10^{-7}$ | 0.11 | $9 \times 10^{-6}$ | -0.012 | 0.2 |
| 19:48643050:CCT:C | *LIG1* | 19AM2 | 36.87 | -0.083 | $1.2 \times 10^{-7}$ | -0.057 | 0.0016 | -0.0073 | 0.29 |
| 19:48620943:C:A | *LIG1* | 19AM3 | 0.17 | 1.2 | $5.9 \times 10^{-11}$ | 0.89 | $1.2 \times 10^{-5}$ | -0.17 | 0.022 |

**Supplementary Table 4. Effects of haplotypes influencing age-at-onset and age-at-SDMT30 for Huntington's disease on *TCF4* repeat expansion in blood.** For each tag variant previously associated with age-at-onset (z-score) or age-at-SDMT30 (z-score) for Huntington's disease (Table S1 of ref. [39]), the effect size and $p$-value from our meta-analysis of *TCF4* somatic expansion in UKB+AoU are shown. The effect allele is the ALT allele for all but one variant; for the 5AM1 variant for which the effect allele is the REF allele, it is typeset in bold and underlined.

| | | | | Somatic *HTT* CAG expansion in blood (ref. [41]) | | Somatic *TCF4* CAG expansion in blood (UKB+AoU) | |
|---|---|---|---|---|---|---|---|
| Locus | Tag variant | Modifier haplotype | MAF (%) | Beta | $p$-value | Beta | $p$-value |
| *MSH2,MSH6* | chr2:47491330:T:C | 2ABEM1 | 1.4 | -0.23 | $3.7 \times 10^{-42}$ | -0.00 | 0.96 |
| *MSH2,MSH6* | chr2:47978940:**T**:G | 2ABEM2 | 37.0 | 0.03 | $1.3 \times 10^{-9}$ | -0.02 | 0.0097 |
| *HTT* | chr4:3074723:C:T | 4ABEM1 | 3.4 | 0.13 | $4.8 \times 10^{-30}$ | -0.02 | 0.35 |
| *MSH3* | chr5:80632699:T:C | 5ABEM1 | 24.4 | 0.04 | $7.2 \times 10^{-19}$ | -0.08 | $6.9 \times 10^{-25}$ |
| *MSH3* | chr5:80660180:T:G | 5ABEM3 | 24.5 | -0.04 | $4.6 \times 10^{-21}$ | 0.05 | $2.7 \times 10^{-12}$ |
| *MSH3* | chr5:80790685:A:G | 5ABEM2 | 0.4 | -0.22 | $1.2 \times 10^{-10}$ | 0.01 | 0.79 |
| *PMS2* | chr7:5978813:A:G | 7ABEM3 | 4.4 | 0.05 | $8.4 \times 10^{-8}$ | 0.05 | 0.0018 |
| *PMS2* | chr7:5986899:C:T | 7ABEM1 | 1.8 | 0.10 | $9.4 \times 10^{-11}$ | 0.05 | 0.021 |
| *PMS2* | chr7:6006003:T:C | 7ABEM4 | 1.0 | 0.09 | $1.9 \times 10^{-6}$ | 0.08 | 0.0022 |
| *PMS2* | chr7:6021277:**A**:G | 7ABEM5 | 18.0 | 0.03 | $1.5 \times 10^{-7}$ | 0.02 | 0.058 |
| *PMS2* | chr7:6063110:A:G | 7ABEM2 | 6.4 | 0.05 | $3.4 \times 10^{-8}$ | 0.03 | 0.023 |
| *MLH3* | chr14:75017975:**GTC**:G | 14ABEM1 | 46.8 | -0.03 | $4.1 \times 10^{-10}$ | 0.02 | 0.00043 |
| *MLH3* | chr14:75017978:**T**:A | 14ABEM1 | 46.8 | -0.03 | $4.1 \times 10^{-10}$ | 0.02 | 0.00043 |
| *FAN1* | chr15:30873651:**C**:T | 15ABEM2 | 44.2 | 0.05 | $3.3 \times 10^{-39}$ | 0.06 | $3.9 \times 10^{-23}$ |
| *FAN1* | chr15:30905792:C:T | 15ABEM3 | 0.8 | 0.15 | $4.9 \times 10^{-12}$ | 0.16 | $1 \times 10^{-5}$ |
| *FAN1* | chr15:30910758:G:A | 15ABEM1 | 0.9 | 0.18 | $2.5 \times 10^{-18}$ | 0.20 | $6.6 \times 10^{-11}$ |
| *FAN1* | chr15:30912434:C:T | 15ABEM4 | 1.8 | 0.07 | $6.4 \times 10^{-6}$ | 0.10 | $1.6 \times 10^{-5}$ |
| *ATAD5, ADAP2,TEFM* | chr17:30836573:T:A | 17ABEM1 | 4.3 | 0.06 | $4.9 \times 10^{-9}$ | 0.07 | $1.8 \times 10^{-5}$ |
| *ATAD5, ADAP2,TEFM* | chr17:30936682:C:T | 17ABEM2 | 22.3 | 0.03 | $8.5 \times 10^{-8}$ | 0.04 | $2.2 \times 10^{-7}$ |

**Supplementary Table 5. Effects of haplotypes influencing *HTT* repeat expansion in blood on *TCF4* repeat expansion in blood.** For each tag variant previously associated with somatic expansion of the *HTT* repeat in blood DNA (Table S2 of ref. [41]), the effect size and $p$-value from our meta-analysis of *TCF4* somatic expansion in UKB+AoU are shown. The effect allele is the minor allele, which is the ALT allele for most variants; for variants for which the effect (minor) allele is the REF allele, it is typeset in bold and underlined. Effect sizes (betas) for somatic expansion of *HTT* are in units of the adjusted somatic expansion ratio (SER) defined in ref. [41]; for *TCF4*, units of betas are standard deviations of our *TCF4* somatic-expansion phenotype. The scatter plot of betas shown in Fig. 4C is restricted to variants that reached $p < 5 \times 10^{-8}$ in the GWAS of *HTT* expansion in blood [41].

| Locus | Category | Quantitative trait | $\beta$ (s.e.) | $p$-value |
|---|---|---|---|---|
| *GLS* | Anthropometric | Height | -0.47 (0.09) | $7.3 \times 10^{-8}$ |
| *GLS* | Blood | Lymphocyte Ct. | -0.45 (0.09) | $3.7 \times 10^{-7}$ |
| *GLS* | Bone and joint | Alkaline Phosphatase | 0.56 (0.09) | $9.4 \times 10^{-10}$ |
| *GLS* | Bone and joint | Calcium | 0.44 (0.09) | $3.5 \times 10^{-6}$ |
| *GLS* | Liver | Gamma Glutamyltransferase | 0.71 (0.09) | $6.8 \times 10^{-15}$ |
| *GLS* | Liver | Aspartate Aminotransferase | 0.42 (0.09) | $4.8 \times 10^{-6}$ |
| *GLS* | Renal | Cystatin C | 0.45 (0.09) | $6.9 \times 10^{-7}$ |
| *TCF4* | Anthropometric | BMI | -0.04 (0.005) | $3.3 \times 10^{-12}$ |
| *TCF4* | Blood | White Ct. | -0.04 (0.005) | $6.9 \times 10^{-13}$ |
| *TCF4* | Blood | Lymphocyte Ct. | -0.04 (0.005) | $1.1 \times 10^{-11}$ |
| *TCF4* | Blood | Neutrophil Ct. | -0.03 (0.005) | $4 \times 10^{-8}$ |
| *TCF4* | Blood | Platelet Ct. | -0.03 (0.005) | $4.6 \times 10^{-6}$ |
| *TCF4* | Blood pressure | Diastolic BP | 0.03 (0.006) | $1.1 \times 10^{-6}$ |
| *TCF4* | Blood pressure | Systolic BP | 0.02 (0.006) | $1.3 \times 10^{-5}$ |
| *TCF4* | Cancer | Testosterone | -0.05 (0.008) | $5.7 \times 10^{-9}$ |
| *TCF4* | Cancer | IGF1 | 0.03 (0.006) | $1.6 \times 10^{-6}$ |
| *TCF4* | Diabetes | HbA1c | 0.03 (0.006) | $4 \times 10^{-10}$ |
| *TCF4* | Liver | Gamma Glutamyltransferase | -0.03 (0.006) | $1.1 \times 10^{-6}$ |
| *TCF4* | Lung | FEV1/FVC | 0.03 (0.006) | $2.4 \times 10^{-8}$ |
| *TCF4* | Renal | Creatinine | -0.04 (0.006) | $6.2 \times 10^{-13}$ |
| *TCF4* | Renal | Urate | -0.03 (0.006) | $1.8 \times 10^{-9}$ |
| *TCF4* | Renal | Cystatin C | -0.03 (0.006) | $1.8 \times 10^{-6}$ |
| *TCF4* | Renal | Urea | -0.02 (0.006) | $3.6 \times 10^{-5}$ |
| *DMPK* | Blood | Hemoglobin | 0.32 (0.05) | $2.6 \times 10^{-10}$ |
| *DMPK* | Blood | Red Ct. | 0.27 (0.05) | $1.1 \times 10^{-7}$ |
| *DMPK* | Blood | Immature Reticulocyte Fraction | 0.22 (0.05) | $3.2 \times 10^{-5}$ |
| *DMPK* | Blood | High Light Scatter Reticulocyte Ct. | 0.22 (0.05) | $3.3 \times 10^{-5}$ |
| *DMPK* | Liver | Gamma Glutamyltransferase | 0.24 (0.05) | $5.3 \times 10^{-6}$ |
| *DMPK* | Lung | FVC | -0.23 (0.06) | $2.9 \times 10^{-5}$ |
| *DMPK* | Renal | Cystatin C | 0.62 (0.05) | $1 \times 10^{-32}$ |
| *DMPK* | Renal | Total Protein | -0.25 (0.05) | $3.5 \times 10^{-6}$ |

**Supplementary Table 6. Associations of CAG repeat expansions with heritable quantitative traits.** Associations that reached a significance threshold of $p < 5 \times 10^{-5}$ (slightly conservative for testing long-allele carrier status at 14 CAG repeat loci for association with 57 quantitative traits) are listed. For biomarker traits, category names (Bone and joint, Cancer, Diabetes, Renal, Liver) are from UK Biobank's biomarker panel documentation. Effect sizes are in units of standard deviations.

| Locus | Disease phenotype | Odds ratio | 95% CI | $p$-value |
|---|---|---|---|---|
| *HTT* | G10 (Huntington's disease) | 11250 | 1914-66100 | $5.5 \times 10^{-25}$ |
| *TCF4* | H18 (other disorders of cornea) | 2.97 | 2.71-3.25 | $3.7 \times 10^{-123}$ |
| *DMPK* | G47 (sleep disorders) | 2.64 | 1.8-3.87 | $7.3 \times 10^{-7}$ |
| *DMPK* | G71 (primary disorders of muscles) | 396.9 | 246-640.5 | $1.1 \times 10^{-132}$ |
| *DMPK* | H25 (senile cataract) | 2.74 | 2-3.75 | $3.6 \times 10^{-10}$ |
| *DMPK* | H26 (other cataract) | 5.89 | 4.65-7.47 | $9.2 \times 10^{-49}$ |
| *DMPK* | H28 (cataract and other disorders of lens in diseases classified elsewhere) | 142.2 | 33.6-601.5 | $1.6 \times 10^{-11}$ |
| *DMPK* | I42 (cardiomyopathy) | 5.65 | 2.87-11.13 | $5.3 \times 10^{-7}$ |
| *DMPK* | I44 (atrioventricular and left bundle-branch block) | 4.08 | 2.67-6.24 | $7.5 \times 10^{-11}$ |

**Supplementary Table 7. Associations of CAG repeat expansions with disease phenotypes.** Associations that reached a significance threshold of $p < 10^{-6}$ (conservative for testing long-allele carrier status at 14 CAG repeat loci for association with 1,129 "first occurrence" binary disease phenotypes) are listed.

| Category | Quantitative trait | *GLS* pLoF SNP/indel $\beta$ (s.e.) | *GLS* pLoF SNP/indel $p$-value | Long but not highly expanded *GLS* $\beta$ (s.e.) | Long but not highly expanded *GLS* $p$-value | Highly expanded *GLS* $\beta$ (s.e.) | Highly expanded *GLS* $p$-value |
|---|---|---|---|---|---|---|---|
| Anthropometric | Height | -0.29 (0.098) | 0.0029 | -0.44 (0.16) | 0.0068 | -0.47 (0.1) | $6.3 \times 10^{-6}$ |
| Blood | Lymphocyte Ct. | -0.14 (0.099) | 0.15 | -0.31 (0.17) | 0.065 | -0.51 (0.11) | $2.4 \times 10^{-6}$ |
| Blood | Platelet Distr. Width | 0.021 (0.098) | 0.83 | -0.05 (0.17) | 0.76 | 0.46 (0.11) | $2 \times 10^{-5}$ |
| Blood | Platelet Ct. | -0.032 (0.099) | 0.75 | -0.23 (0.17) | 0.17 | -0.38 (0.11) | 0.00042 |
| Bone and joint | Alkaline Phosphatase | 0.058 (0.1) | 0.56 | 0.18 (0.17) | 0.29 | 0.71 (0.11) | $1.2 \times 10^{-10}$ |
| Bone and joint | Calcium | -0.046 (0.1) | 0.65 | 0.14 (0.18) | 0.45 | 0.56 (0.11) | $6.3 \times 10^{-7}$ |
| Bone and joint | Vitamin D | 0.017 (0.1) | 0.87 | -0.077 (0.17) | 0.66 | -0.39 (0.11) | 0.00052 |
| Cancer | IGF1 | -0.14 (0.1) | 0.15 | 0.15 (0.17) | 0.38 | -0.38 (0.11) | 0.0006 |
| Liver | GGT | -0.083 (0.1) | 0.41 | 0.15 (0.17) | 0.37 | 0.96 (0.11) | $3.5 \times 10^{-18}$ |
| Liver | AST | -0.0071 (0.1) | 0.94 | 0.26 (0.17) | 0.13 | 0.5 (0.11) | $7.5 \times 10^{-6}$ |
| Liver | Albumin | -0.085 (0.1) | 0.41 | 0.43 (0.18) | 0.019 | -0.41 (0.11) | 0.00022 |
| Renal | Cystatin C | 0.027 (0.1) | 0.79 | 0.02 (0.17) | 0.91 | 0.59 (0.11) | $7.8 \times 10^{-8}$ |
| Renal | Phosphate | 0.058 (0.1) | 0.57 | -0.01 (0.18) | 0.95 | -0.45 (0.11) | $7.7 \times 10^{-5}$ |
| | Glutamine | 0.44 (0.13) | 0.001 | 0.71 (0.22) | 0.0011 | 0.78 (0.14) | $1.9 \times 10^{-8}$ |

**Supplementary Table 8. Associations of highly expanded *GLS* repeats with heritable quantitative traits.** Bonferroni-significant associations ($p < 0.05/57$, adjusting for 57 traits tested) for highly expanded *GLS* repeat status (i.e., ≥1 IRR pair) are shown. Effect sizes and $p$-values for association tests of less-expanded *GLS* repeat status (i.e., anchored IRRs at *GLS*, but no IRR pairs) and *GLS* SNP/indel pLoF status are also provided. Effect sizes and $p$-values are from a joint model of pLoF status and expansion status (including both highly expanded and less-expanded repeat status as independent regressors), adjusting for sex, age, and age squared. Association results are also shown for glutamine. GGT, gamma-glutamyl transferase; AST, aspartate aminotransferase.

| | UK Biobank | | | |
|---|---|---|---|---|
| Phenotype | | OR | 95% CI | $p$-value |
| K75 | Other inflammatory liver diseases | 9.8 | 2.0-47.1 | 0.0043 |
| K75.4 | Autoimmune hepatitis | 34.9 | 3.8-319.4 | 0.0017 |
| K76 | Other diseases of liver | 3.6 | 1.4-8.8 | 0.0061 |
| K76.8 | Other specified diseases of liver | 8.2 | 2.2-31.2 | 0.0019 |
| N18 | Chronic kidney disease (CKD) | 4.5 | 2.3-9.0 | $1.6 \times 10^{-5}$ |
| N18.5 | Chronic kidney disease, stage 5 | 23.7 | 6.2-90.5 | $3.5 \times 10^{-6}$ |
| N18.9 | Chronic kidney disease, unspecified | 5.3 | 2.2-12.8 | 0.00023 |
| N18.0 or N18.5 | CKD stage 5, inclusive of end-stage renal disease | 20.6 | 5.5-76.8 | $7.0 \times 10^{-6}$ |
| | *All of Us* | | | |
| Other diseases of liver | | 2.3 | 0.8-6.7 | 0.12 |
| Chronic kidney disease (CKD) | | 4.8 | 1.9-12.3 | $1.2 \times 10^{-3}$ |
| CKD stage 5, inclusive of end-stage renal disease | | 10.1 | 3.0-34.2 | $1.9 \times 10^{-4}$ |
| | Meta-analysis | | | |
| Other diseases of liver | | 3.0 | 1.5-5.9 | $2.0 \times 10^{-3}$ |
| Chronic kidney disease (CKD) | | 4.6 | 2.6-8.1 | $7.0 \times 10^{-8}$ |
| CKD stage 5, inclusive of end-stage renal disease | | 14.0 | 5.7-34.3 | $7.2 \times 10^{-9}$ |

**Supplementary Table 9. Associations of highly expanded *GLS* repeats with liver and kidney diseases.** Associations that reached FDR<0.05 with top-level ICD-10 categories (3 significant out of 11 categories tested, K70–K77 and N17–N19) and associations that reached FDR<0.05 with ICD-10 subcategories (4 significant out of 62 subcategories tested) are listed. Upon closer examination of ICD-10 subcategory definitions within N18, we determined that combining N18.5 (chronic kidney disease, stage 5) and N18.0 (end-stage renal disease) would be a better choice for a CKD stage 5 phenotype, so we ran an association test for this combined phenotype in UKB and took this phenotype forward (along with two category-level phenotypes) for replication in AoU. Finally, we meta-analyzed the UKB and AoU associations (log odds ratios) using a linear random-effects model (DerSimonian-Laird estimator) using the rma.uni() function within the metafor package in R.
